# Inhibition of Renin Release, a Crucial Event in Homeostasis, is Mediated by Coordinated Calcium Oscillations within Juxtaglomerular Cell Clusters

**DOI:** 10.1101/2024.12.23.629519

**Authors:** Hiroki Yamaguchi, Nick A. Guagliardo, Laura A. Bell, Manako Yamaguchi, Daisuke Matsuoka, Fang Xu, Jason P. Smith, Mohamed Diagne, Lucas F. Almeida, Silvia Medrano, Paula Q. Barrett, Edward H. Nieh, R. Ariel Gomez, Maria Luisa S. Sequeira-Lopez

## Abstract

**BACKGROUND:** Juxtaglomerular (JG) cells are sensors that control blood pressure (BP) and fluid-electrolyte homeostasis. They are arranged as clusters at the tip of each afferent arteriole. In response to a decrease in BP or extracellular fluid volume, JG cells secrete renin, initiating an enzymatic cascade that culminates in the production of angiotensin II (AngII), a potent vasoconstrictor that restores BP and fluid-electrolyte homeostasis. In turn, AngII exerts negative feedback on renin release concomitantly with increased intracellular Ca^2+^, preventing excessive circulating renin and hypertension. However, within their native structural organization, the intricacies of intracellular Ca^2+^ signaling dynamics and their sources remain uncharacterized.

**METHODS:** We generated mice expressing the JG cell-specific genetically encoded Ca^2+^ indicator (GCaMP6f) to investigate Ca^2+^ dynamics within JG cell clusters *ex vivo* and *in vivo*. For *ex vivo* Ca^2+^ imaging, acutely prepared kidney slices were perfused continuously with a buffer containing variable Ca^2+^ and AngII concentrations ± Ca^2+^ channel inhibitors. For *in vivo* Ca^2+^ image capture, native mouse kidneys were imaged *in situ* using multi-photon microscopy with and without AngII administration. ELISA measurements of renin concentrations determined acute renin secretion *ex vivo* and *in vivo*, respectively.

**RESULTS:** *Ex vivo* Ca^2+^ imaging revealed that JG cells exhibit robust and coordinated intracellular oscillatory signals with cell-cell propagation following AngII stimulation. AngII dose-dependently induced stereotypical burst patterns characterized by consecutive Ca^2+^ spikes, which inversely correlated with renin secretion. Pharmacological channel inhibition identified key sources of these oscillations: endoplasmic reticulum Ca^2+^ storage and release, extracellular Ca^2+^ uptake via ORAI channels, and intercellular communication through gap junctions. Blocking ORAI channels and gap junctions reduced AngII inhibitory effect on renin secretion. *In vivo* Ca^2+^ imaging demonstrated robust intracellular and intercellular Ca^2+^ oscillations within JG cell clusters under physiological conditions, exhibiting spike patterns consistent with those measured in *ex vivo* preparations. Administration of AngII enhanced the Ca^2+^ oscillatory signals and suppressed acute renin secretion *in vivo*.

**CONCLUSION:** AngII elicits coordinated intracellular and intercellular Ca^2+^ oscillations within JG cell clusters, *ex vivo* and *in vivo*. The effect is driven by endoplasmic reticulum-derived Ca^2+^ release, ORAI channels, and gap junctions, leading to suppressed renin secretion.

## INTRODUCTION

Precise inhibition of renin release is crucial in the regulation of blood pressure (BP) and fluid-electrolyte homeostasis. Renin, the key regulated hormone of the renin-angiotensin system (RAS), is secreted by juxtaglomerular (JG) cells that are arranged in clusters of 7–10 cells at the tip of the afferent arterioles near the glomeruli. Within such clusters, JG cells communicate with one another and other components of the juxtaglomerular apparatus (JGA)^1^, including the afferent and efferent arterioles, the extraglomerular mesangium, and the macula densa^2^. This unique microenvironment and complex anatomical arrangement allow JG cells to act as sensitive detectors of small changes in perfusion pressure and the composition and volume of the extracellular fluid^2,3^. Thus, the aforementioned interactions of renin cells within the JGA ensure that renin release and its suppression are finely adjusted to sustain homeostasis.

Intracellular Ca^2+^ [Ca^2+^]_int_ acts as a proximal regulator of renin synthesis and secretion in JG cells and is influenced by various physiological stimuli^4–6^. Ca^2+^ exhibits a paradoxical relationship with renin. In contrast to most other secretory cells where Ca^2+^ stimulates hormone production (e.g., zona-glomerulosa cells of adrenal^7^), [Ca^2+^]_int_ elevation in JG cells suppresses renin production and release. [Ca^2+^]_int_ suppresses renin synthesis and secretion in JG cells by seemingly inhibiting cyclic adenosine monophosphate (cAMP), a potent stimulator of renin synthesis and secretion^5,6^ and a determinant of renin cell identity via its epigenomic profile^8,9^. Despite its crucial importance in renin regulation, the dynamics of [Ca^2+^]_int_ and how it is regulated in JG cells remain poorly understood.

Renin initiates the RAS cascade that culminates in the production of angiotensin II (AngII), the primary effector of the system. In turn, AngII suppresses renin production concomitant with an elevation in [Ca^2+^]_int_ levels in JG cells. While cell culture studies have confirmed these relationships^5^, *in vivo* evidence of this direct negative feedback is lacking^10^. In 1989, Kurtz et al. reported that AngII induces [Ca^2+^]_int_ oscillatory patterns in isolated mouse JG cells^11^. However, such finding was limited by the absence of the complex cellular interactions present in the native kidney structure. As indicated above, JG cells depend highly on their surrounding tissue environment for proper function and safeguard of their identity. Further, they are tightly coupled by gap junctions (GJs), which facilitate cell-to-cell passage of signaling molecules and inorganic ions among coupled JG cells, JGA components, vascular smooth muscle cells (VSMCs), and endothelial cells^2,6,12,13^. In fact, pure JG cells isolated from kidney tissue devoid of the connections mentioned above, stop synthesizing renin within 72 hours and differentiate into VSMCs^2,14^. The scarcity of JG cells, their intricate organization and location, and plasticity of JG cells have hindered studies on Ca^2+^ dynamics within their native structure and environment.

Using a novel mouse model expressing a genetically encoded Ca^2+^ indicator targeted to JG cells, we elucidated [Ca^2+^]_int_ dynamics in JG cells and determined the source of [Ca^2+^]_int_ oscillations and the channel mediators involved in *ex vivo* and *in vivo* renin release within a native structural and physiological context.

## METHODS

### Data Availability

The detailed methods and Major Resources Table are available in Supplemental Material. The corresponding authors can provide data supporting the study’s findings upon request.

## RESULTS

### AngII evokes periodic Ca^2+^ signals within the JG cell cluster (JGCC) in the *ex vivo* kidney

To study the [Ca^2+^]_int_ dynamics in JG cells, we crossed *Ren1^c-Cre/+^* mice, where Cre expression is driven by the *Ren1^c^* promoter^15^, with floxed-*GCaMP6f* mice^16^, generating mice with JG cell-specific GCaMP6f expression (*Ren1^c^-GCaMP6f* mice; Figure 1A). *Ren1^c^-GCaMP6f* mice showed no overt renal phenotype, as evidenced by their unaltered kidney vascular development, plasma renin levels, and *Ren1* mRNA expression (Figure S1). First, we confirmed JG cell-specific GCaMP6f expression in *Ren1^c^-GCaMP6f* mice by immunofluorescence labeling of renin in the kidney cortex and colocalization with GCaMP6f. Renin-positive cells are highly colocalized with GCaMP6f expression in cells located within the JGA (Figure 1B, Figure S2). To measure acute changes in [Ca^2+^]_int_ elicited by AngII in JG cells, we prepared kidney slices (75 μm-thick) from *Ren1^c^-GCaMP6f* mice, capturing GCaMP6 fluorescence with a water-immersion objective lens under continuous perfusion with PIPES buffer (Figure 1C). Without AngII, JG cells were quiescent, with minimal Ca^2+^ activity. In contrast, 3 nM AngII elicited periodic blinking patterns of [Ca^2+^]_int_ in distinct cells within each JGCC (Figure 1D, Video S1). Upon closer inspection of JGCCs, changes in fluorescence propagated through the cytoplasm (Ca^2+^ wave) successively between adjacent JG cells (Figure 1E). A representative kymograph plot indicates that lines tracking time-dependent Ca^2+^ activity of distinct JG cells and the sequential Ca^2+^ activity between adjacent cells, suggesting cell-cell interaction among the JGCC members. Furthermore, to quantify the Ca^2+^ dynamics in distinct JG cells, we employed the imaging analysis platform Mesmerize^17^ and CaImAn^18^ toolset for motion correction, unbiased region of interest (ROI) detection, and signal demixing/extraction of each time series (Figure 1F). This approach allowed us to quantify the number of Ca^2+^ spikes elicited by AngII in distinct JG cells within JGCCs, highlighting the robust [Ca^2+^]_int_ oscillatory activity. Collectively, our *ex vivo* imaging of kidney slices from *Ren1^c^-GCaMP6f* mice enabled us to characterize spatial-temporal Ca^2+^ dynamics in JG cells, demonstrating that AngII elicits robust Ca^2+^ oscillations within JGCCs.

**Figure 1.**
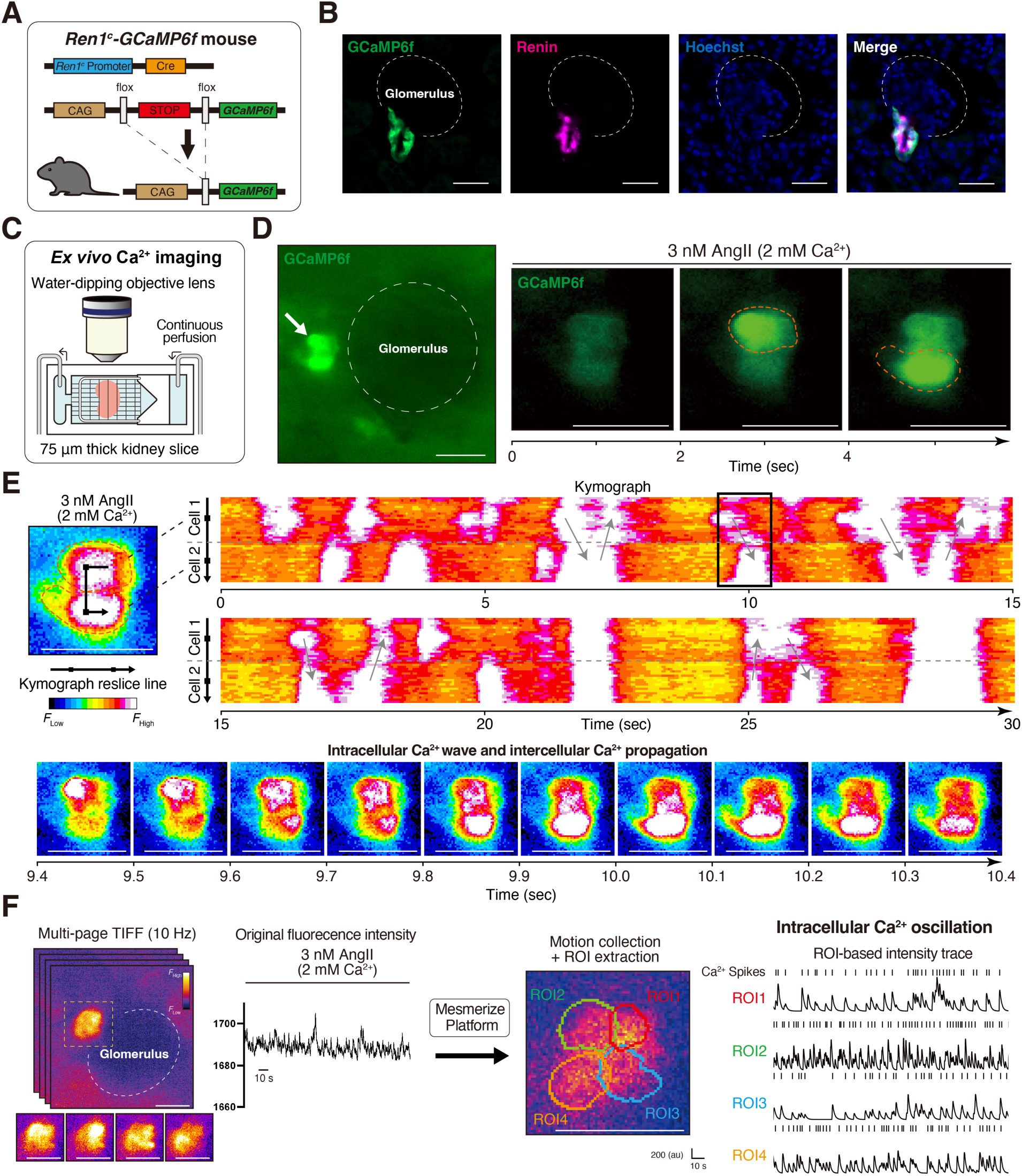
Angiotensin II (AngII) elicits robust and periodic Ca^2+^ oscillations propagating across juxtaglomerular cell clusters (JGCCs) expressing GCaMP6f in *ex vivo* kidneys. **A,** Schematic of *Ren1^c^-GCaMP6f* mouse. **B,** Immunofluorescence staining for renin in the kidney cortex of the *Ren1^c^-GCaMP6f* mice. The expression of GCaMP6f is colocalized with renin in the JG area. See also Figure S2. **C,** Schematic of *ex vivo* Ca^2+^ imaging using acutely prepared kidney slices from *Ren1^c^-GCaMP6f* mice. **D**, Time-series of captured images perfused with 3 nM AngII in 2 mM Ca^2+^-containing buffer. White arrows point to GCaMP6f-expressing JG cells. Orange-dashed lines encircle the individual JG cells adjacent to each other. Individual JG cells can be distinguished by cytosolic GCaMP6f blinking periodically (see also Video S1). **E,** Time-series of fluorescent intensity of GCaMP6f-expressing JG cells shown by the kymograph (top), indicating changes in Ca^2+^ activity across designated reslice lines (black arrows) spanning adjacent JG cells (gray arrows). Gray-dotted lines indicate the border of adjacent JG cells. Images at 0.1-second intervals (bottom) designated by a black square in the kymograph visualize oscillatory Ca^2+^ waves in distinct JG cells and the sequential Ca^2+^ activity between adjacent cells. **F,** Schematic strategy for *ex vivo* Ca^2+^ spike analysis. Mesmerize platform was employed to perform motion correction and region of interest (ROI) extraction from spatiotemporal cropped JGCCs, exhibiting Ca^2+^ transient spikes in each ROI with raster plots corresponding to the intensity trace peaks. All scale bar, 20 μm.

### AngII elicits Ca^2+^ spike bursts that correlate with renin secretion

Next, we characterized the pattern and dose-dependency of the Ca^2+^ activity elicited by AngII in JGCCs. In kidney slices treated with one of four doses of AngII for 25 minutes, Ang II elicited a dose-dependent increase in Ca^2+^ spikes (Figure 2A). Spike intervals exhibited an unimodal distribution, featuring a distinct peak around 2.0–2.5 seconds, with no variations across the AngII concentrations (Figure 2B). Notably, these spikes were evoked in “bursts” defined as three or more consecutive spikes with inter-spike periods shorter than the specified threshold (event periods, Figure S3A, S3B)^7^. While the mean event periods were consistent across the AngII concentrations (Figure 2C), the mean number of bursts/min within JGCCs was AngII dose-dependent (Figure 2D). Conversely, the mean burst durations and the number of Ca^2+^ spikes per burst remained invariant among AngII doses (Figure 2E). Furthermore, the dose-dependence of evoked Ca^2+^ bursts was inversely correlated with the dose-dependency of renin suppression (Figure 2F, S4A–S4B). Thus, in aggregate, our data support Ca^2+^ spike bursts as a driver of renin inhibition.

**Figure 2.**
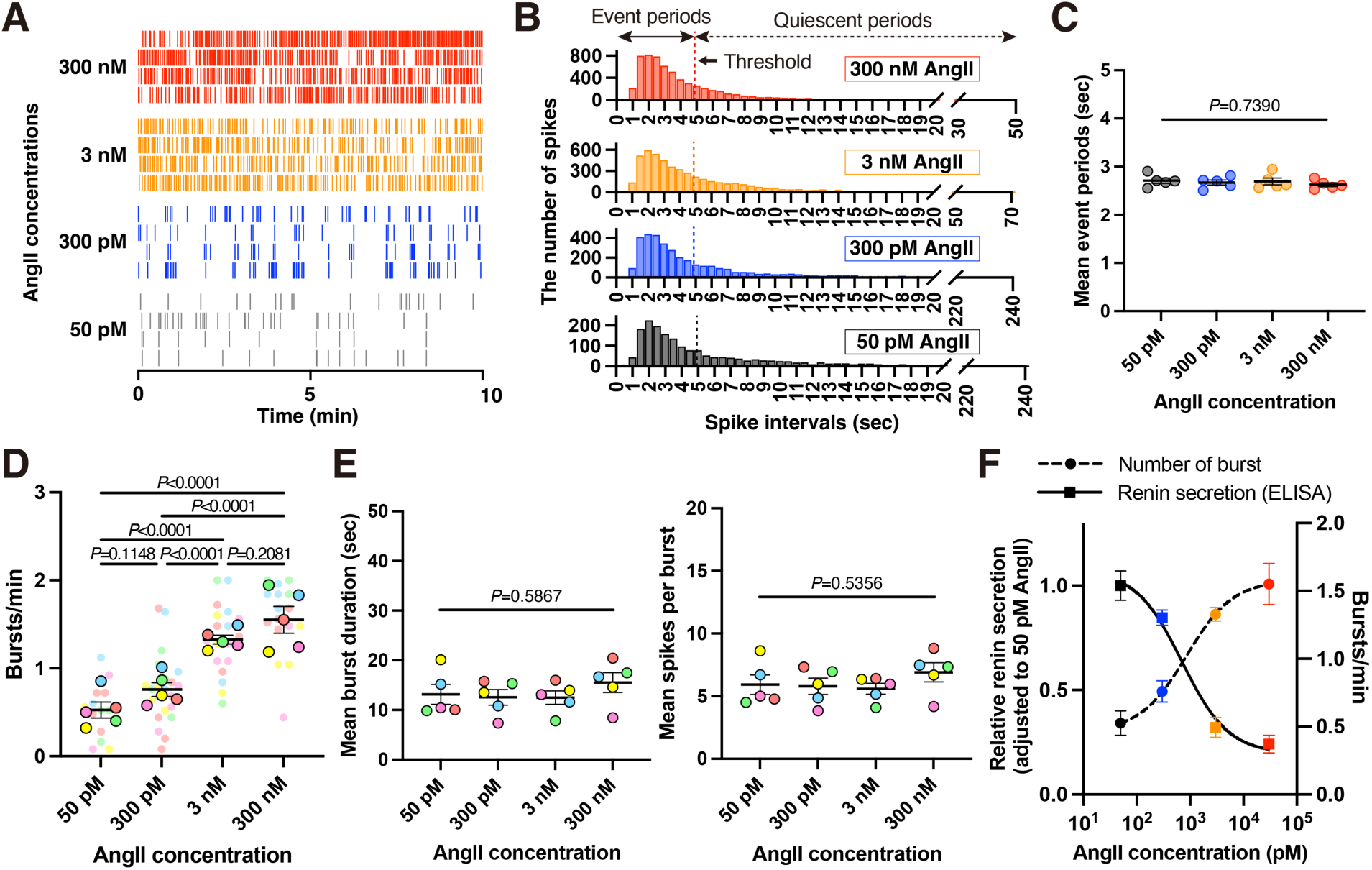
Angiotensin II (AngII) dose-dependently elicits stereotypical Ca^2+^ spike patterns and suppresses renin secretion within juxtaglomerular cell clusters (JGCCs). **A,** Representative raster plots for Ca^2+^ spikes in JGCCs under treatment with 50 pM, 300 pM, 3 nM, and 300 nM AngII. **B–E,** Analysis for Ca^2+^ spikes in JGCCs of *Ren1^c^-GCaMP6f* mice imaged over 25 minutes under treatment with 50 pM (n=4 mice, 5 slices, 15 ROIs), 300 pM (n=5 mice, 5 slices, 22 ROIs), 3 nM (n=5 mice, 5 slices, 18 ROIs), and 300 nM AngII (n=5 mice, 5 slices, 17 ROIs). Each slice contains one JGCC analyzed. **B,** Histogram of spike intervals across all ROIs exhibit unimodal distribution across AngII concentrations. The threshold distinguishes between event periods (active phase comprising bursts) and quiescent periods (inactive inter-burst phase). See also Figure S3. **C,** Mean event periods in JGCCs showed no notable variances across AngII concentrations (One-way ANOVA). **D,** The number of bursts/min is dose-dependently increased, especially between 300 pM and 3 nM AngII (linear mixed model with Tukey’s multiple comparison test). Identical color circles represent ROIs from the same JGCCs. **E,** Mean burst duration (left) and spike per burst (right) showed no significant difference across AngII concentrations (One-way ANOVA). **F,** The fold increase in renin secretion in kidney slice preparations after stimulation with AngII concentrations, plotted relative to unstimulated secretion and adjusted to the value of 50 pM AngII (left y-axis, n=4 slices per concentration, 2-month-old male C57BL/6J mice, non-linear regression fit: R^2^=0.92, EC_50_=707.1 pM). The adjusted renin secretion is inversely correlated with the dose-dependence of the number of bursts/min per slice (right y-axis, non-linear regression fit: R^2^=0.81, EC_50_=860.0 pM). Data presented as mean±SEM.

### JG cells import extracellular Ca^2+^ mainly through ORAI channels to mediate AngII-elicited Ca^2+^ oscillations and suppress renin secretion

Next, we asked if the AngII-elicited Ca^2+^ spikes require Ca^2+^ from extracellular sources. Perfusion of 3 nM AngII with the Ca^2+^ chelator, ethyl-glycol tetraacetic acid (EGTA), in a “Ca^2+^-free” buffer (0.1 mM Ca^2+^) nearly eliminated Ca^2+^ oscillations (96.33% decrease), while restoring Ca^2+^ in the perfusion buffer (Recovery) induced renewed Ca^2+^ activity, indicating extracellular Ca^2+^ is necessary for Ca^2+^ oscillations in JG cells (Figure 3A). To confirm whether changes in Ca^2+^ activity in Ca^2+^-free buffer parallel functional changes in renin secretion, we measured renin from kidney slices in response to AngII with and without extracellular Ca^2+^. Kidney slices treated with 3 nM AngII for 30 min markedly decreased renin secretion in the presence of 2 mM Ca^2+^ but did not elicit a change in renin in the Ca^2+^-free condition (Figure 3B, S4B). Combined, our *ex vivo* Ca^2+^ imaging analysis and renin secretion assays highlight the dependence of robust oscillatory Ca^2+^ signals on extracellular Ca^2+^.

**Figure 3.**
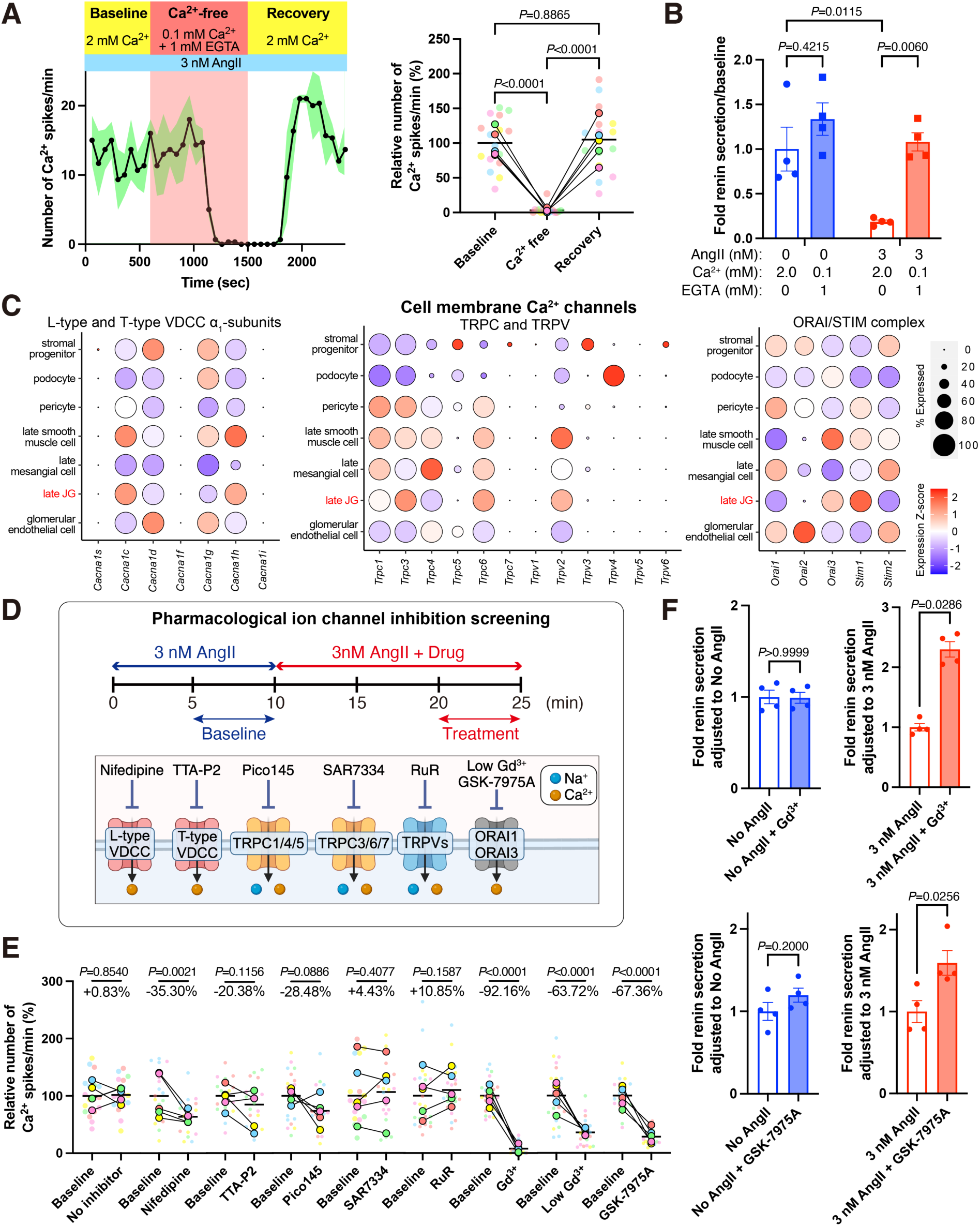
Angiotensin II (AngII)-elicited Ca^2+^ oscillations and renin secretion in juxtaglomerular cell clusters (JGCCs) depend on extracellular Ca^2+^, primarily via ORAI channels. **A,** A representative time-course of Ca^2+^ spikes/min in each ROI (n=3) within *Ren1^c^-GCaMP6f* mouse kidney slice (left). The green-shaded areas represent SEM. The right panel summarizes the effect of transiently removing and washing back extracellular Ca^2+^ on JGCCs (adjusted to Baseline, n=4 mice, 5 slices, n=16 ROIs, linear mixed model with Tukey’s multiple comparison test). **B,** The result of kidney slice renin ELISA with 2 mM Ca^2+^ and 0.1 mM Ca^2+^ + 1 mM ethyl-glycol tetraacetic acid (EGTA) buffer under No-AngII and 3 nM AngII (n=4 per concentration, two-way ANOVA; Šídák’s multiple comparison test). **C.** Bubble plots for gene expression profiles of L- and T-type voltage-dependent Ca^2+^ channels (VDCC) α_1_-subunits, TRPC and TRPV subfamilies, and ORAI/STIM complex in the renin lineage cell-related clusters. **D,** Schematic for pharmacological ion channel inhibition screening. The timeline (top) and target channels with inhibitors (bottom) are illustrated. **E,** The relative number of Ca^2+^ spikes/min (%) in captured JG cells adjusted to Baseline (No-inhibitor, n=5 mice, 5 slices, 18 ROIs; 10 μM Nifedipine, n=4 mice, 5 slices, 18 ROIs; 10 μM TTA-P2, n=5 mice, 5 slices, 18 ROIs; 10 μM Pico145, n=5 mice, 5 slices, 17 ROIs; 10 μM SAR7334, n=4 mice, 5 slices, 20 ROIs; 10 μM Ruthenium red (RuR), n=4 mice, 5 slices, 25 ROIs; 50 μM Gd^3+^, n=4 mice, 5 slices, 23 ROIs; 5 μM Gd^3+^ (Low Gd^3+^), n=4 mice, 5 slices, 22 ROIs; 10 μM GSK-7975A, n=4 mice, 5 slices, 20 ROIs, linear mixed model). Identical color circles represent ROIs from the same JGCCs. **F,** The result of kidney slice renin ELISA treated with 50 μM Gd^3+^ (top) and 10 μM GSK-7975A (bottom) under 3 nM and No-AngII (n=4 per concentration, 2-month-old male C57BL/6J mice, Mann-Whitney U test). Data presented as mean±SEM.

We then investigated the cell membrane Ca^2+^ conductances responsible for maintaining AngII-elicited Ca^2+^ oscillations in JG cells. To identify candidate Ca^2+^ channels, we analyzed the gene expression profiles of major cytoplasmic Ca^2+^ permeable channels using single-cell RNA-seq data of mouse kidney cells derived from *FoxD1^+^* precursors, which include renin lineage cells, isolated as previously reported^9^ (Figure 3C). This analysis revealed the high expression of the following genes in the late JG cluster: i). Ca_v_1.2 (*Cacna1c*), Ca_v_1.3 (*Cacna1d*), Ca_v_3.1 (*Cacna1g*), and Ca_v_3.2 (*Cacna1h*), α_1_-subunits of L-type and T-type voltage-dependent Ca^2+^ channels (VDCC); ii). TRPC1, TRPC3, TRPC4, TRPC6, among transient receptor potential channel canonical family (TRPC) and TRPV2 in TRP vanilloid family (TRPV); and iii). ORAI1 and ORAI3, the pore-forming subunits of store-operated Ca^2+^ channels (SOCC) and their Ca^2+^-sensors, stromal interaction molecule (STIM) proteins^19^. Based on these findings, we screened candidate Ca^2+^ conductances using pharmacological channel inhibitors and *ex vivo* Ca^2+^ imaging (Figure 3D). For each drug, JGCCs were imaged for 10 minutes with 3 nM AngII perfusion (Baseline), followed by the addition of channel inhibitors for 15 minutes (Treatment), and mean Ca^2+^ spikes/minute in the last 5 minutes of baseline and treatment conditions were compared. Applying 3 nM AngII maintained consistent Ca^2+^ activity throughout 25-minute imaging experiments; prolonged exposure to 488 nm fluorescence alone did not significantly influence the Ca^2+^ response to AngII (Figure 3E).

#### L-type and T-type VDCCs

L-type and T-type VDCC are crucial in generating and maintaining Ca^2+^ oscillations in various cell types, including endocrine cells^7^. L-type VDCC exhibits high conductance with slow inactivation upon membrane depolarization^20^ and is reported to be functionally expressed in isolated JG cells^21^, while T-type channels are low conductance Ca^2+^ channels activated by small membrane depolarizations and are rapidly inactivated^20^. Dihydropyridine nifedipine (a potent Ca_v_1 antagonist) moderately decreased AngII-elicited Ca^2+^ spikes (35.30% decrease), and TTA-P2^22^ (a pan Ca_v_3 family antagonist) did so partially (20.38% decrease), yet neither fully eliminated the Ca^2+^ spikes (Figure 3E).

#### TRP channels

Based on the reports that the TRPC-mediated pathway inhibits renin release from JG cells^23^, we targeted TRPC1/4/5 and TRPC3/6/7 subfamilies, which form non-selective, Ca^2+^-permeable hetero-multimeric cation channels^24^ with Pico145^25^ and SAR7334^26^, respectively. In addition, we used ruthenium red (RuR), a pan TRPV inhibitor^27^, to target TRPV2 as well as TRPV4, which was reported to function as a mechanosensitive channel in renin cells^28^. Inhibition with Pico145, SAR7334, or RuR only partially or did not reduce AngII-elicited Ca^2+^ spikes (28.48% decrease, 4.43% increase, and 10.85% increase, respectively; Figure 3E).

#### ORAI channels

To assess the contribution of ORAI channels to JG cell Ca^2+^ oscillation, we blocked both ORAI1 and ORAI3 channels. Treatment with 50 μM trivalent lanthanide gadolinium (Gd^3+^), a potent inhibitor of all ORAI isoforms (ORAI1, ORAI2, and ORAI3)^29,30^, drastically inhibited the number of Ca^2+^ spikes (92.16% decrease, Figure 3E) elicited by AngII. In addition, Gd^3+^ prevented the inhibition of renin secretion in 30 minutes under 3 nM AngII but not in the absence of AngII (Figure 3F). Since Gd^3+^ is also reported to affect other cell membrane cation channels^31^, we applied more selective ORAI inhibitors: low concentrations of Gd^3+^ (5 μM), which functions as an inhibitor of all ORAI isoforms^29,30^ with greater selectivity, and pyrazole derivatives GSK-7975A, which inhibits recombinant ORAI1 and ORAI3 currents^32^. Both 5 μM Gd^3+^ and GSK-7975A significantly suppressed the number of Ca^2+^ spikes induced by 3 nM AngII (5 μM Gd^3+^; 63.72% decrease, GSK-7975A; 67.36% decrease, Figure 3E). Furthermore, GSK-7975A significantly attenuated the inhibition of renin secretion in kidney slices stimulated with 3 nM AngII but not in the absence of AngII (Figure 3F).

Combined, this data indicates that ORAI channels contribute to maintaining AngII-elicited Ca^2+^-oscillations and the concurrent suppression of renin, while individual VDCC and TRPC1/4/5 channels partially account for Ca^2+^ oscillatory activity in JG cells.

### AngII-elicited Ca^2+^ spikes depend on endoplasmic reticulum (ER) Ca^2+^ storage in JG cells

Given that ORAI/STIM-mediated store-operated Ca^2+^ entry (SOCE) helps maintain [Ca^2+^]_int_ activity by transporting extracellular Ca^2+^ into the ER to facilitate the Ca^2+^ release from the ER^30,33^, we next investigated whether Ca^2+^ release from the ER contributes to AngII-elicited Ca^2+^ oscillations in JG cells. Our single-cell RNA-seq data revealed high expression of sarcoplasmic/endoplasmic reticulum Ca^2+^-ATPase (SERCA) and inositol 1,4,5-trisphosphate (IP3) receptor (IP3R) genes in the late JG clusters, while gene expression of ryanodine receptors (RyR) was low (Figure 4A). Thus, we examined SERCA, which transports cytosolic Ca^2+^ into the ER lumen, thereby controlling the amount of Ca^2+^ in the ER (Figure 4B). If AngII-elicited Ca^2+^ oscillations depend on ER-Ca^2+^ cycling, blocking Ca^2+^ storage should prevent Ca^2+^ spike generation. Indeed, the pharmacological blockade of SERCA with cyclopiazonic acid (CPA)^34^ severely reduced Ca^2+^ spike generation (93.66% decrease, Figure 4C). Furthermore, commensurate with reduced Ca^2+^ spiking, CPA induced a persistent and global increase in fluorescent intensity (Figure 4D), indicating sustained [Ca^2+^]_int_ elevation. This phenomenon is typical of enhanced SOCE via store-operated Ca^2+^ channels (SOCC) triggered by the depletion of ER Ca^2+^ stores^35^.

**Figure 4.**
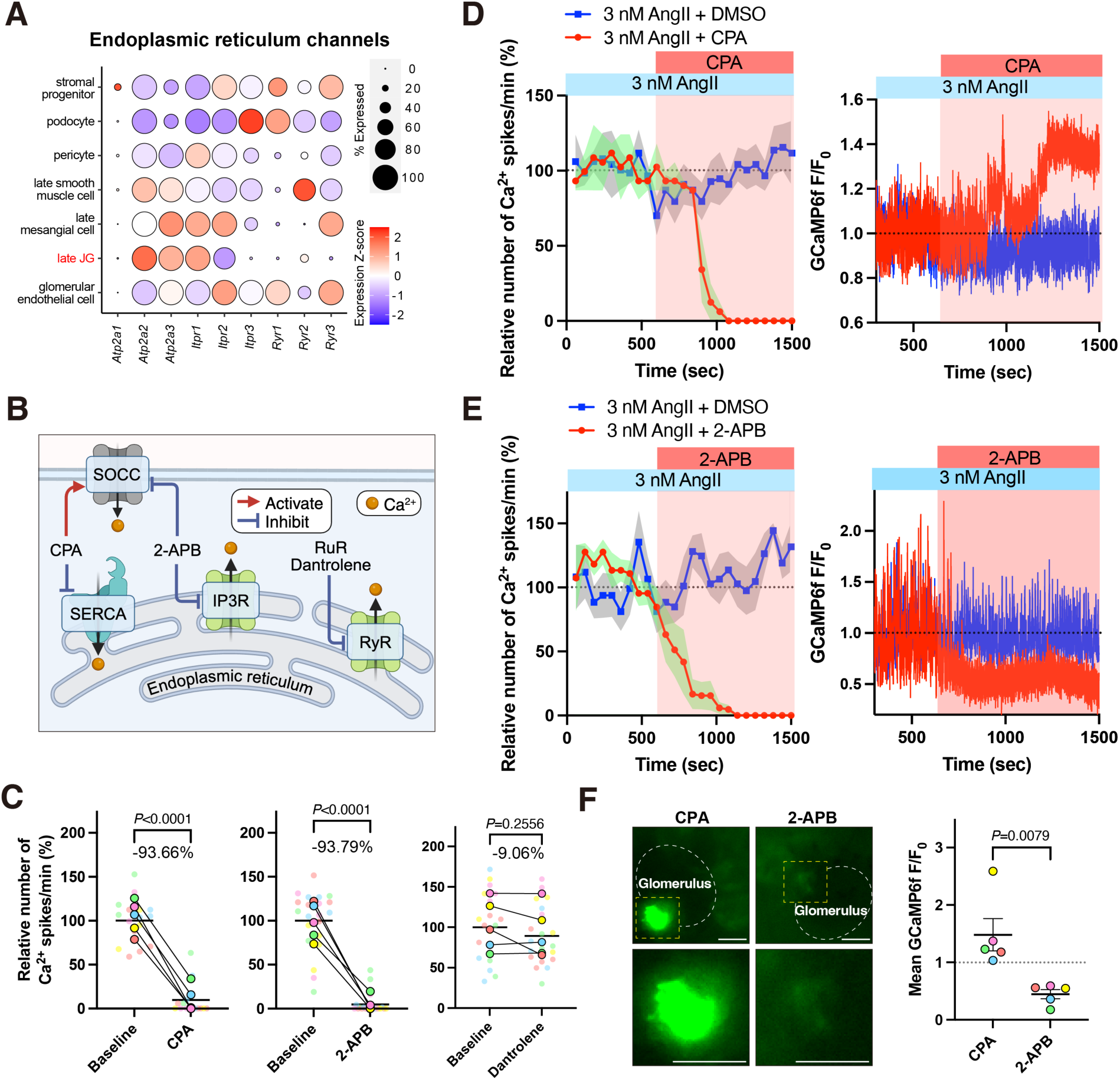
Endoplasmic reticulum (ER) Ca^2+^ stores and releases generate angiotensin II (AngII)-induced Ca^2+^ oscillatory signals in juxtaglomerular cell clusters (JGCCs). **A**, Bubble plots for gene expression profiles of ER channels indicate high gene expression of sarcoplasmic/endoplasmic reticulum Ca^2+^-ATPase (SERCA, *Atp2a*) and inositol 1,4,5-trisphosphate (IP3) receptor (IP3R, *Ip3r*) genes in the JG clusters while showing low gene expression of ryanodine receptors (RyR, *Ryr*). **B,** Schematic for pharmacological ER channel inhibition and the inhibitors. SOCC; store-operated Ca^2+^ channels. **C,** The relative number of Ca^2+^ spikes/min (%) in captured JGCCs adjusted to Baseline (20 μM Cyclopiazonic acid (CPA), n=4 mice, 5 slices, 16 ROIs; 100 μM 2-Aminoethoxydiphenyl borate (2-APB), n=5 mice, 5 slices, 21 ROIs, 10 μM Dantrolene, n=5 mice, 5 slices, 20 ROIs, linear mixed model). Identical color circles represent ROIs from the same JGCCs. **D–E,** Representative time-courses of the number of Ca^2+^ spikes (left) and cytosolic fluorescence intensities (right) before and after perfusion of CPA (**D**, n=2 ROIs) or 2-APB (**E**, n=4 ROIs) under 3 nM AngII treatment. The data for treatment with the same volume of DMSO are shown as controls. The green- and gray-shaded areas represent SEM. **F,** A representative image (left) and a mean GCaMP6f F/F_0_ in Treatment periods (right) with CPA and 2-APB. The GCaMP6f intensities in JGCCs increased in the kidney slice treated with CPA, while those in 2-APB decreased (Mann-Whitney U test). Scale bar, 20 μm. Data presented as mean±SEM.

To examine the dependence of oscillatory activity on other ER release channels, we blocked IP3R with a cell-permeable IP3R inhibitor, 2-Aminoethoxydiphenyl borate (2-APB)^36^, and RyR with RyR1 and RyR3 inhibitor dantrolene^37^. Treatment with 2-APB markedly decreased the Ca^2+^ spikes (93.79% decrease), while dantrolene failed to reduce the number of Ca^2+^ spikes (9.06% decrease, Figure 4C). TRPV inhibitor RuR, which also acts as a RyR inhibitor^38^, further supports the non-contribution of RyRs to AngII-elicited Ca^2+^ oscillations in JG cells (Figure 3I). In contrast to SERCA blockade, IP3R inhibition with 2-APB significantly decreased overall fluorescent intensity while simultaneously decreasing Ca^2+^ spikes (Figure 4F), likely due to 2-APB’s effect as an SOCC inhibitor^39^. This data indicates that AngII-elicited Ca^2+^ oscillations in JG cells depend on Ca^2+^ storage and release from the ER dependent on SERCA and IP3R activities.

### JGCCs require GJs to induce robust AngII-elicited Ca^2+^ oscillations and cell-cell coordination to suppress renin secretion

Our *ex vivo* Ca^2+^ imaging uncovered a time-dependent activation among neighboring JG cells (Figure 1E), raising the possibility of cell-cell communication among the JGCC members. In agreement with previous data demonstrating that GJs are abundantly and functionally expressed in JG cells^12,13,40^, we used our single-cell RNA-seq data to examine the diversity of GJ components in JG cells. We observed high expression of connexin (Cxs) in JG cells and adjacent VSMCs and endothelial cells (e.g., Cx37, Cx40, Cx43, and Cx45; Figure 5A). To investigate the contribution of GJs to Ca^2+^ activity among neighboring JG cells, we applied a pan GJ inhibitor, carbenoxolone (CBX)^41^, and analyzed the Ca^2+^ signal pattern within the JGCCs. Notably, CBX effectively prevented AngII-induced Ca^2+^ propagation across adjacent JG cells within the clusters (Figure 5B, Video S2) while markedly reducing the number of Ca^2+^ spikes (94.18% decrease, Figure 5C). Finally, we confirmed that CBX restored renin secretion within 30 minutes after 3 nM AngII exposure but failed to alter secretion in its absence (Figure 5D). These data support the requirement for GJs involvement in the mechanism whereby Ca^2+^ oscillatory activity is evoked by AngII to suppress renin secretion.

**Figure 5.**
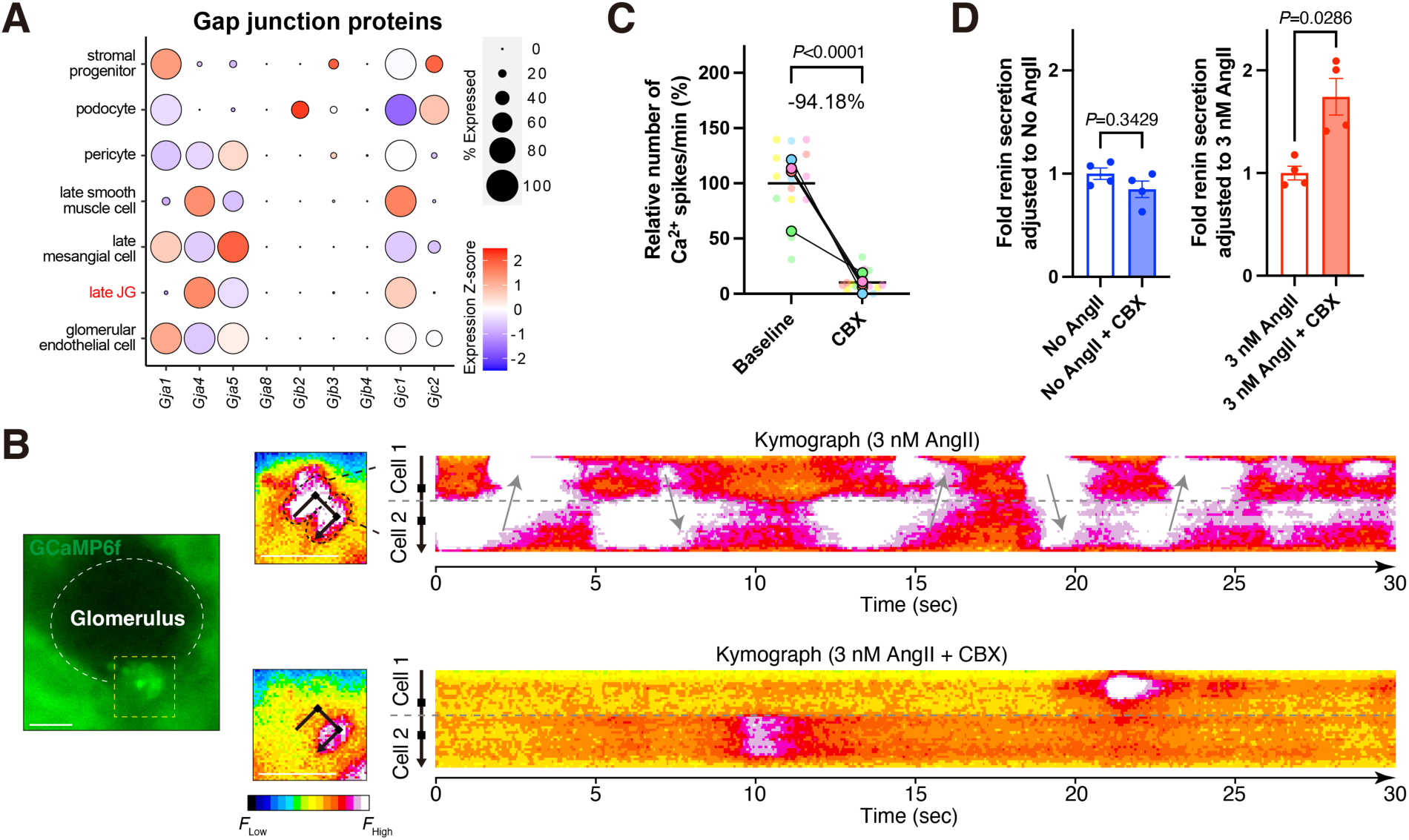
Gap junction (GJ) inhibition suppresses intracellular and intercellular Ca^2+^ activity within the juxtaglomerular cell clusters (JGCCs). **A,** Bubble plots for gene expression profiles of GJ indicate diverse and highly expressed gene expression profiles of GJ proteins, including Connexin43 (Cx43, *Gja1*), Cx37 (*Gja4*), Cx40 (*Gja5*), and Cx45 (*Gjc1*) in JG cells and their adjacent cells. **B,** Representative kymographs show the time-series of fluorescent intensity under 3 nM angiotensin II (AngII) and 3 nM AngII + 100 μM carbenoxolone (CBX) perfusion using *Ren1^c^-GCaMP6f* mouse kidney slice. Black arrows designate reslice lines. Black-dashed lines encircle the individual JG cells adjacent to each other. Gray arrows indicate the spanning Ca^2+^ activity across the border of adjacent JG cells (gray-dashed lines). See also Video S2. Scale bar, 20 μm. **C**, The relative number of Ca^2+^ spikes/min (%) in captured JGCCs under 3 nM AngII treatment with 100 μM CBX (n=4 mice, 5 slices, n=16 ROIs, linear mixed model). Identical color circles represent ROIs from the same JGCCs. **D,** The kidney slice renin ELISA results with or without 100 μM CBX under 3 nM and no AngII (n=4 per concentration, 2-month-old male C57BL/6J mice, Mann-Whitney U test). Data presented as mean±SEM.

### *In vivo* JGCCs actively generate intracellular and intercellular Ca^2+^ oscillations under physiological conditions

To explore the Ca^2+^ dynamics within native JGCCs *in vivo*, we performed intravital imaging of intact living *Ren1^c^-GCaMP6f* mouse kidneys using multi-photon microscopy (Figure 6A). First, we assessed *in vivo* Ca^2+^ signals within the JGCCs under physiological conditions. High-speed continuous volume images (29.989 Hz, captured of 81 μm volume z-stack) detected robust oscillatory Ca^2+^ signals in the JG cells that propagated within the JGCCs (Figure 6B, Video S3). To characterize *in vivo* Ca^2+^ spike patterns in distinct JG cells, we applied the same strategy as *ex vivo* Ca^2+^ spike analysis to continuous t-series images captured over 3 minutes (Figure 6C). Consequently, the spike intervals of *in vivo* JGCCs displayed an unimodal distribution with a distinct peak around 2.0–2.5 seconds, demonstrating spike-interval distribution patterns and mean event periods similar to *ex vivo* treatment with AngII (Figure 6D). In addition, the mean duration of bursts and the number of Ca^2+^ spikes within a burst remained unchanged between *in vivo* and *ex vivo* samples treated with each AngII dosage (Figure 6E). The number of spikes and bursts per minute *in vivo* was nearly identical to those in *ex vivo* kidneys treated with 3 nM or 300 nM AngII (Figure 6F). Collectively, our *in vivo* Ca^2+^ imaging of *Ren1^c^-GCaMP6f* mouse kidneys revealed that native JGCCs actively generate intracellular and intercellular Ca^2+^ oscillations, similar to those observed in *ex vivo* Ca^2+^ imaging treated with high-dose AngII.

**Figure 6.**
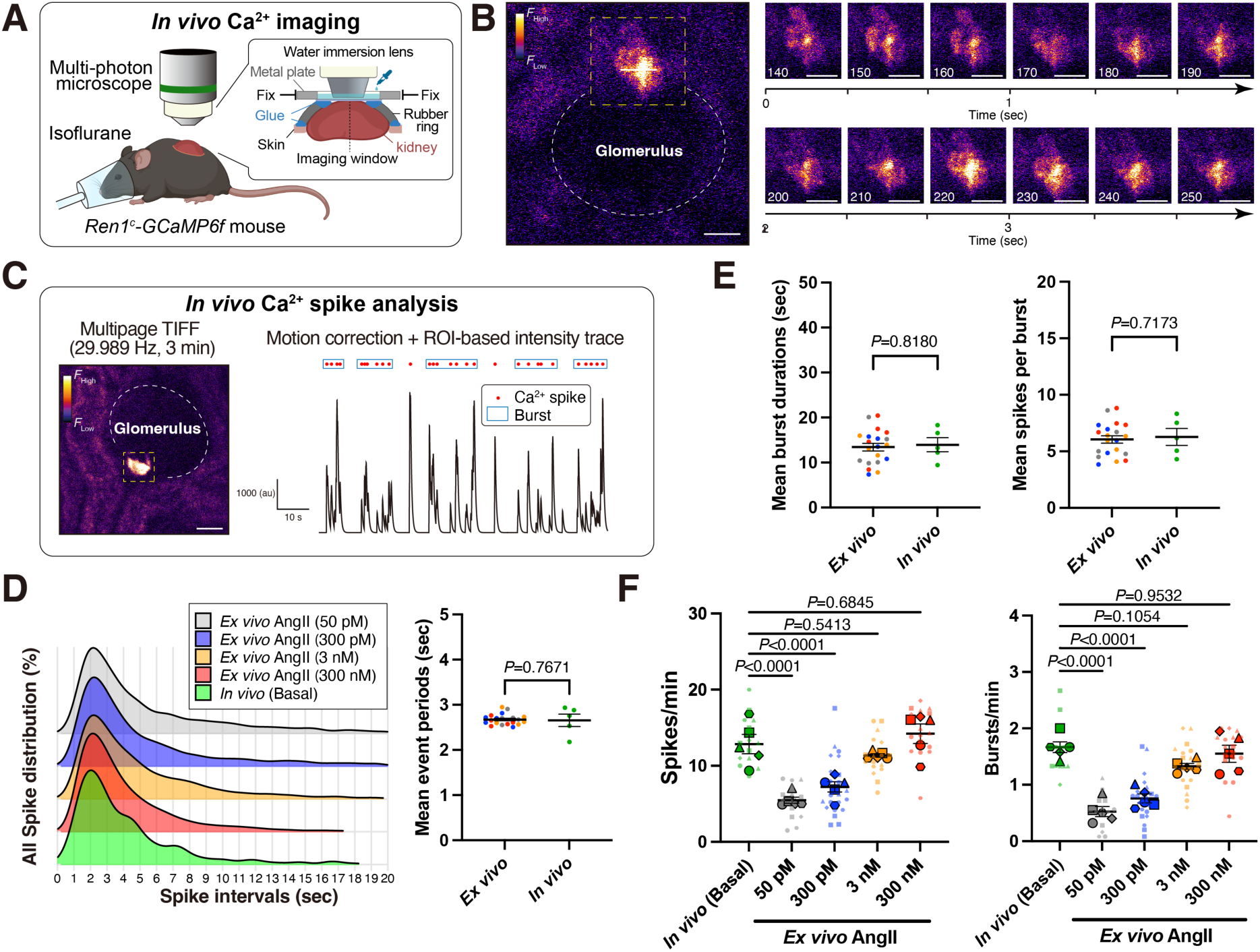
*In vivo* juxtaglomerular cell clusters (JGCCs) generate coordinated intracellular and intercellular Ca^2+^ oscillations within the native clusters. **A,** Schematic of *in vivo* Ca^2+^ imaging of JGCCs in *Ren1^c^-GCaMP6f* mice. **B,** Captured fluorescence volume series of GCaMP6f exhibit robust oscillatory signals propagate across the JGCCs within the cluster. (29.989 Hz). A yellow-dashed square line indicates a JGCC, shown on the right in representative images at 10-image intervals. See also Video S3. **C,** Schematic strategy for *in vivo* Ca^2+^ spike analysis. Multi-page TIFF images captured in 3 minutes were used to trace the fluorescence intensity for the spatiotemporal JGCCs, following the *ex vivo* Ca^2+^ imaging strategy shown in Figure 1F. **D**, The ridgeline plots (left) for Ca^2+^ spikes patterns across all ROIs in five JGCCs captured by *in vivo* Ca^2+^ imaging under basal state (green, n=4 mice, 5 JGCCs, 17 ROIs) and *ex vivo* Ca^2+^ imaging treated with each AngII concentration (gray; 50 pM, blue; 300 pM, orange; 3 nM, red; 300 nM. Mean event periods (right) showed no significant difference between *in vivo* Ca^2+^ imaging and *ex vivo* in all AngII treated groups (Mann-Whitney U test). **E,** Mean burst duration (left) and spike per burst (right) showed no significant difference between *in vivo* Ca^2+^ imaging and *ex vivo* for all AngII treated groups combined (Mann-Whitney U test). **F,** The number of spikes/minute (left) and bursts/minute (right) indicate that those observed *in vivo* under basal conditions were nearly identical to those in *ex vivo* treated with 3 nM or 300 nM AngII (linear mixed model with Tukey’s multiple comparison test). Identical shape symbols represent ROIs from the same *in vivo* kidney or kidney slice. All scale bar, 20 μm. Data presented as mean±SEM.

### AngII administration enhances Ca^2+^ bursts within the JGCC and suppresses acute renin secretion *in vivo*

Although *in vitro* studies demonstrated that AngII directly inhibits renin secretion concurrent with increased [Ca^2+^]_int_ levels^5^, *in vivo* evidence of the direct effect on JG cells remained controversial^10^. To determine if the physiological Ca^2+^ oscillation within the JGCCs *in vivo* depends on AngII, we continuously captured the signals of identical JGCCs in *Ren1^c^-GCaMP6f* mouse kidneys using multi-photon microscopy during systemic AngII administration. This serial intravital imaging revealed that bolus AngII injection (100 ng/g) through a peritoneal catheter markedly enhanced the Ca^2+^ signalings (Figure 7A, Video S4). The signals began to enhance within 30 seconds of starting AngII administration, exhibiting a substantial burst-like oscillatory signaling of [Ca^2+^]_int_ (Figure 7B). The quantification of GCaMP6f intensity in JGCCs exhibited a significant increase following AngII administration (Figure 7C), corroborating that AngII also enhances the activity of [Ca^2+^]_int_ signaling in native JG cells *in vivo*.

**Figure 7.**
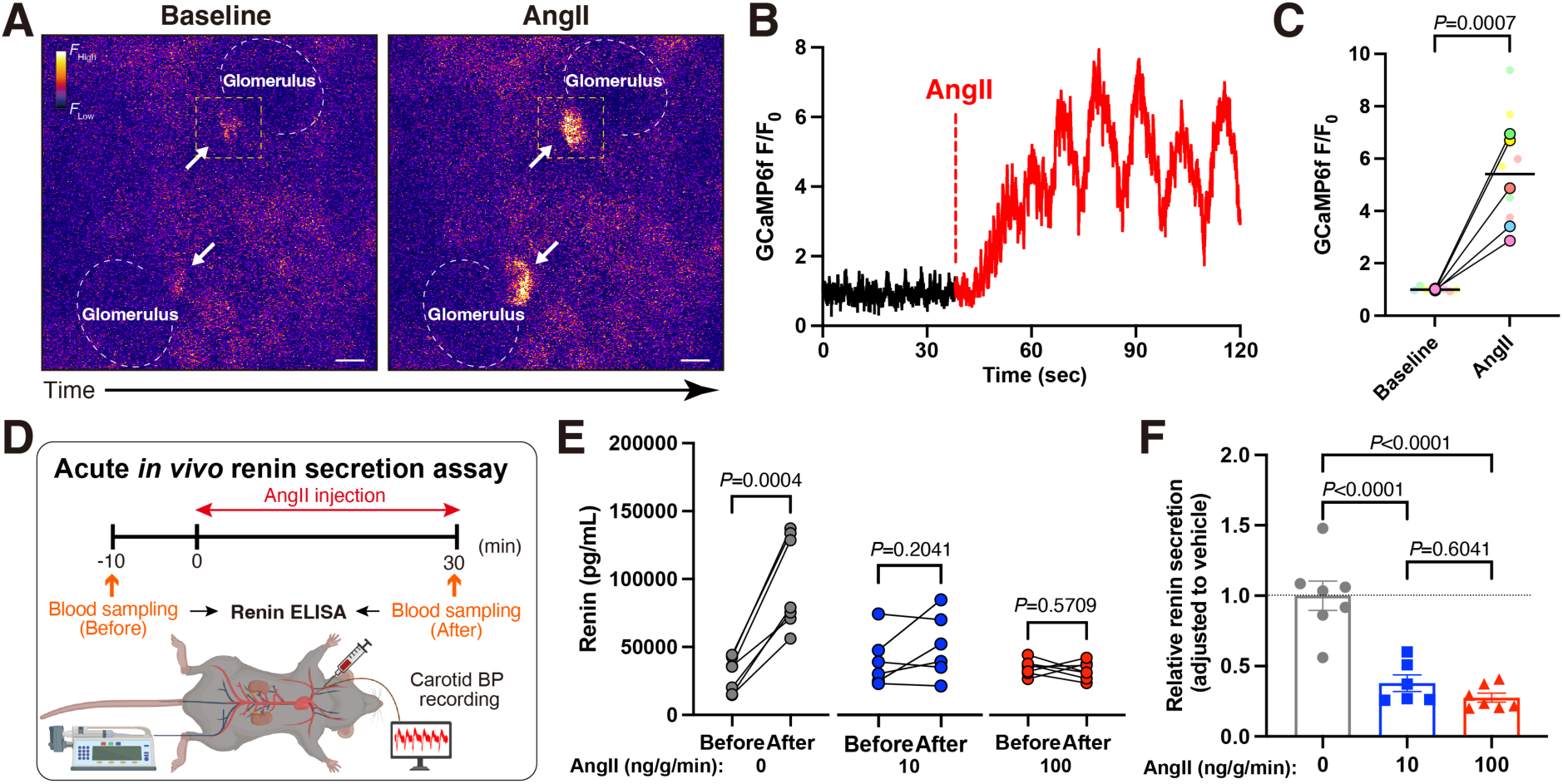
Angiotensin II (AngII) enhances Ca^2+^ oscillatory activity within the native juxtaglomerular cell clusters (JGCCs) and inhibits acute renin secretion *in vivo*. **A–B**, Representative images of GCaMP6f signals from JGCCs in *Ren1^c^-GCaMP6f* mice before (Baseline) and 30 seconds after bolus AngII (100 ng/g) intraperitoneal administration (**A**) and the time-series of GCaMP6f intensities (**B**) in the cropped JGCC (dashed-yellow lines). White arrows represent JGCCs. Scale bar, 20 μm. See also Video 4S. **C**, A mean GCaMP6f intensity was significantly increased 30 seconds after AngII administration (n=5 mice, 8 JGCCs, linear mixed model). Mean GCaMP6f F/F_0_ of 300 multipage TIFFs before and 30 seconds after bolus AngII injection were compared. Identical color circles represent distinct JGCCs captured within the same mouse. **D,** Schematic of acute *in vivo* renin secretion assay. **E–F**, plasma renin levels before and after surgical procedures under continuous administration with 0 (saline vehicle), 10, and 100 ng/g/min AngII (**E,** n=7, 6, and 7 *Ren1^c^-GCaMP6f* mice, respectively, paired t-test) and the ratio adjusted to vehicle (**F**, One-way ANOVA with Tukey’s multi-comparison test). Figure S5 describes carotid blood pressure (BP) data. Data presented as mean±SEM.

Finally, to test whether AngII modulates acute renin secretion *in vivo*, we assessed the change in plasma renin levels before and 30 minutes after the continuous administration of AngII in bolus doses used for *in vivo* Ca^2+^ imaging (100 ng/g/min) and the lower dose (10 ng/g/min) under carotid BP monitoring (Figure 7D). AngII dose-dependently increased carotid BP, while vehicle administration had a minor effect (Figure S5), confirming the effective delivery of Ang II by intraperitoneal administration to the circulation. As expected, and to maintain BP (Figure 7E, gray dots) circulating renin increased 2.5-fold after 30 minutes post blood withdrawal for baseline renin measurement (Figures 7F). AngII administration suppressed such renin secretion and increased BP in a dose-dependent manner (Figure 7E, 7F), indicating that JG cells sensed and responded appropriately to the physiological challenge and its natural inhibitor, AngII. In aggregate, our *in vivo* Ca^2+^ imaging and acute physiological experiments demonstrated that AngII enhances physiological Ca^2+^ oscillations within the JGCCs and acutely inhibits renin secretion *in vivo*.

## DISCUSSION

Cells fluctuate in their cytoplasmic Ca^2+^ concentration to regulate versatile cellular processes — gene transcription, secretion, muscle contraction, development, etc.— and display diverse spatiotemporal Ca^2+^ dynamics^42^. In primary JG cell culture, AngII induces oscillatory [Ca^2+^]_int_ signals that are infrequent (1–2 per minute) and require a supraphysiological concentration of AngII (1 μM)^11^. In other cell types (e.g., epithelial cells), Ca^2+^ signaling patterns in intact tissue are more active than those in isolated cells due to the various signaling mechanisms from surrounding tissues, including Ca^2+^ or IP3 exchanges through GJs, paracrine signals, or Ca^2+^ influx through stretch-activated channels activated by mechanically coupled cells^42,43^. Here, we generated *Ren1^c^-GCaMP6f* mice in which GCaMP expression is highly specific to JG cells within the kidney without affecting renin expression, secretion, or kidney development. This mouse model enabled the study of Ca^2+^ dynamics in JG cells while retaining their native cluster structure *ex vivo* and *in vivo*. In *ex vivo* kidney slices, robust bursts of Ca^2+^ oscillations in JG cells were elicited by physiological concentrations of Ang II propagating among adjacent JG cells, with over ten times higher frequency than previously reported in isolated JG cells^11^. These findings indicate that intercellular interactions within the native kidney structure facilitate Ca^2+^ signaling in JGCCs (Figure 8).

**Figure 8.**
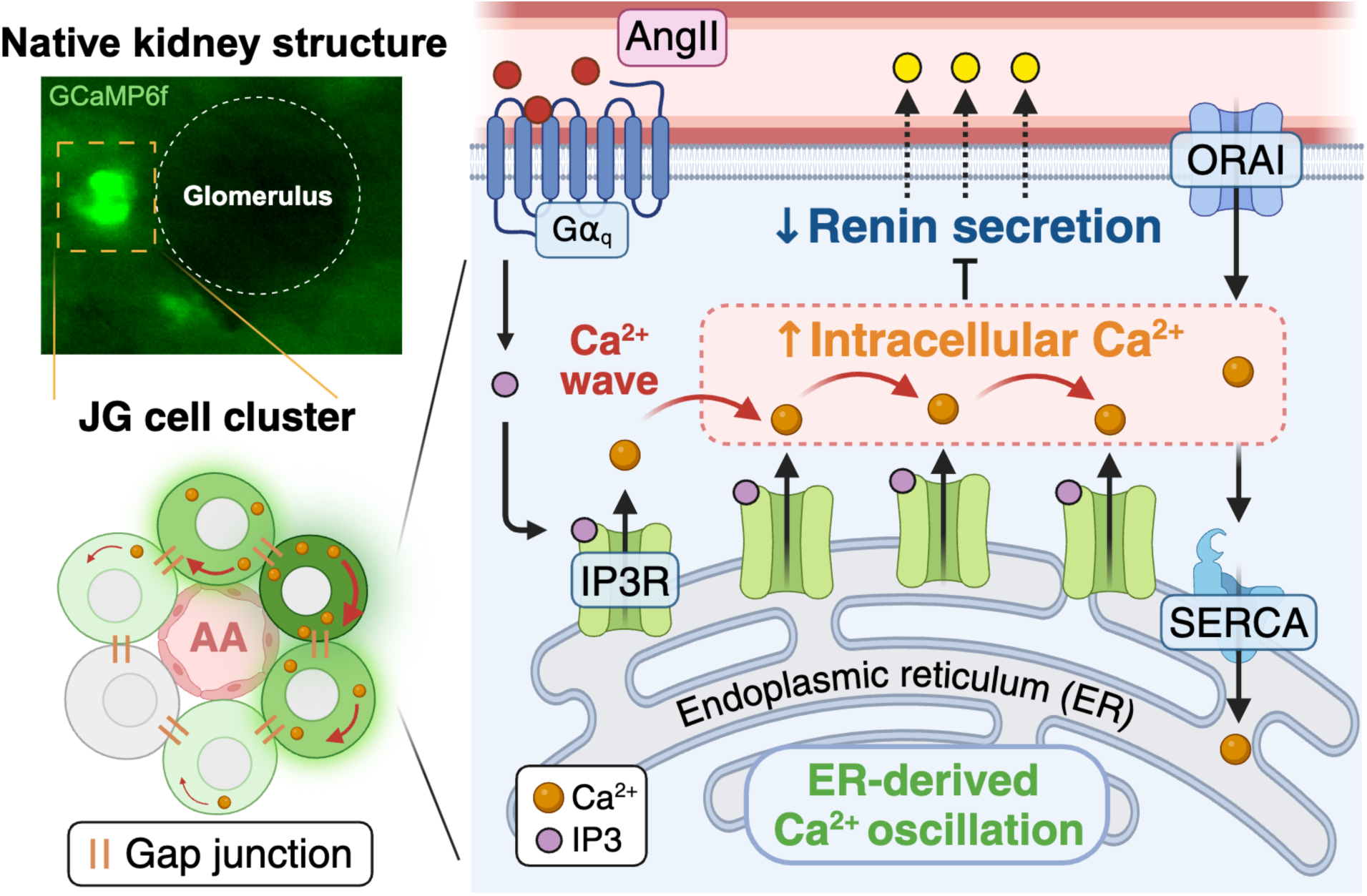
Overview of Angiotensin II (AngII)-elicited Ca^2+^ dynamics within juxtaglomerular cell clusters (JGCCs). AngII induces robust and sustained Ca^2+^-oscillations within their native JGCC structure and suppresses renin secretion. Coordinated Ca^2+^ signaling is mediated by intracellular Ca^2+^ release at the endoplasmic reticulum (ER) supplied through ORAI channels and intercellular gap junctions that propagate signaling molecules. AA; afferent arteriole, IP3R; 1,4,5-trisphosphate (IP3) receptor, SERCA; sarcoplasmic/endoplasmic reticulum Ca^2+^-ATPase.

[Ca^2+^]_int_ concentration is strictly regulated by coordinated crosstalk between multiple ion channels and transporters in the plasma membrane and internal organelles^42^. We investigated the contribution of specific ion channels identified through JG cell’s expression profiling to Ang II-induced [Ca^2+^]_int_ oscillations —and thus renin release— in JG cells by employing a range of pharmacological agents and manipulating extracellular [Ca^2+^] in *ex vivo* kidney slices from *Ren1^c^-GCaMP6f* mice. This approach allowed us to elucidate the complex interplay between ion channels, Ca^2+^ signaling, and renin secretion in the native kidney structure. First, we demonstrate that an extracellular influx of Ca^2+^ is necessary to maintain [Ca^2+^] oscillations; removing Ca^2+^ from the perfusion buffer eliminated AngII-elicited Ca^2+^ spikes. Ca^2+^ activity is reduced, but not eliminated, with an application of the L-type VDCC blocker nifedipine. Unexpectedly, the blockade of other VDCCs and TRP channels reported to be functionally expressed in JG cells^21,23,28^ had a minor effect on AngII-elicited Ca^2+^ spikes, suggesting that the activity of one or more additional ion channels may be required. Next, we demonstrated that [Ca^2+^]_int_ release from intracellular stores is also necessary to maintain AngII-elicited Ca^2+^ oscillations in JG cells. The blockade of IP3R at the ER and the Ca^2+^ uptake inlet SERCA markedly suppressed AngII-elicited Ca^2+^ spikes. In many cell types, AngII binding to the AngII type-1 receptor activates heterotrimeric G-protein (Gα_q_) pathways, inducing phospholipase-C to generate IP3^44^. Both IP3 and Ca^2+^ are required to open IP3R, through which released Ca^2+^ can activate additional, nearby IP3Rs, thus forming the pattern of waves and oscillations^42^. Considering these pathways and our Ca^2+^ imaging data, Ca^2+^ waves and oscillations in JG cells reflect the direct effect of Ca^2+^ release at ER through IP3R (Figure 8).

The presence of SOCCs in JG cells has been inferred from experiments showing that blocking SERCA increases [Ca^2+^]_int_ levels and inhibits renin release in isolated cells^4,5^. Our *ex vivo* Ca^2+^ imaging confirmed that blocking SERCA increased [Ca^2+^]_int_ globally, highlighting the significance of SOCE in JG cells. However, the channels constituting SOCE and their function had not been described in JG cells. In many cell types, STIM/ORAI1 interaction triggers SOCE: STIM proteins at ER sense Ca^2+^ storage depletion and adjust their conformation to activate Ca^2+^ permeable channels composed of ORAI1 at the plasma membrane^19^. Importantly, ORAI/STIM-mediated SOCE contributes to sustaining Ca^2+^ oscillations via IP3R release by transporting extracellular Ca^2+^ into ER, leading to activating transcription factors that regulate gene expression^33^. In particular, ORAI2 and ORAI3, which form hetero-multimers with ORAI1, have recently been identified as crucial for maintaining Ca^2+^ oscillations triggered by physiological receptor activation^30^. Indeed, our pharmacological blockade of ORAI channels targeting both ORAI1 and ORAI3 significantly inhibited Ca^2+^ oscillations and mitigated the suppression of renin secretion induced by AngII in the JGCC (Figure 8). Conditional deletion of ORAI/STIM complexes in cells of the renin lineage *in vivo* would provide profound insights into the role of [Ca^2+^]_int_ signaling in the differentiation and function of renin cells.

Ca^2+^ spreads across groups of cells by coordinating their oscillations, generating synchronous oscillatory Ca^2+^ signals responsible for integrated multicellular behaviors^42^. The mechanisms underlying intercellular Ca^2+^ propagation depend on the diffusion of an intracellular messenger through GJs. Among the GJ proteins, Cxs function as a conduit for intercellular communication by transferring inorganic ions and small signaling molecules, including Ca^2+^, cAMP, and IP3^45^. In mice, the renin cell-specific conditional deletion of Cx40, the most abundant Cxs in JG cells, induces the displacement of renin-expressing cells and inhibits negative regulation of renin synthesis and secretion^40^. However, despite studies showing a critical role for cell-to-cell interactions through GJs for adequate development and function of JG cells, how GJ inhibition affects Ca^2+^ dynamics in JG cells remained uninvestigated. Our *ex vivo* Ca^2+^ imaging suggests that decreased [Ca^2+^]_int_ concentrations may contribute to hyperreninemia in Cx40-deleted animals, as pan-GJ blockade in the JGCCs markedly suppressed [Ca^2+^]_int_ oscillations along with intercellular signaling coordination, mitigating the renin secretion inhibition induced by AngII. These findings suggest that cell-to-cell interactions mediated by GJs are essential for maintaining Ca^2+^ activity within JGCCs, thereby regulating renin secretion (Figure 8). This essential finding could not be detected in conventional cell cultures using isolated JG cells, further highlighting the functional significance of the JG apparatus structure in modulating renin secretion in JG cells.

The intravital imaging of [Ca^2+^]_int_, the crucial proximal regulator for JG cell function, presents several challenges, including the scarcity of JG cells and their anatomical location deep within the blood-rich renal cortex. In this study, we employed a multi-photon microscopy approach to *Ren1^c^-GCaMP6f* mouse kidneys, achieving the capture of [Ca^2+^]_int_ dynamics in native JGCCs *in vivo*. In contrast to *ex vivo* experiments in kidney slices, where JG cells are quiescent in the absence of stimulation, JGCCs *in vivo* produce spontaneous intracellular and intercellular Ca^2+^ oscillations. Further, JG cell Ca^2+^ spikes and bursting activity *in vivo* closely resemble that of *ex vivo* Ca^2+^ activity in AngII-treated kidney slices, suggesting that coordinated Ca^2+^ oscillations are present in the neutral state of native JGCCs within the intact kidney and functional circulatory system. This indicates that, in their neutral state, JGCCs consistently have a brake that retains renin inside the cell, functioning as an effective mechanism for the immediate release of renin when homeostasis is threatened. Additionally, administering AngII to mice resulted in increased Ca^2+^ bursts in native JGCCs and suppressed the acute renin secretion into the circulation (Figure 8). These findings support evidence for AngII’s direct effect on JG cells and the Ca^2+^ oscillation *in vivo*. How the pressure, sympathetic and sensory nerve innervation, and AngII interact in the intact animal to modulate Ca^2+^ dynamics in JG cells remains to be investigated. Intriguingly, mechanosensitive channels that would be inactive *in vitro* may be potent modulators *in vivo*, where the circulatory system maintains physiological pressure.

The role of cAMP in regulating Ca^2+^ dynamics and renin production within the intact kidney is unknown. Due to the ubiquitous expression of cAMP across various cell types —unlike the specific secretion of renin from JG cells— measuring cAMP levels of JG cells in intact kidneys remains a challenge. Since Ca^2+^ dynamics differ in isolated JG cells from those in kidney sections, we predict that cAMP dynamics in JG cells may also be distinct. Simultaneous [Ca^2+^]_int_ and cAMP imaging in JGCCs could provide a more comprehensive understanding of the intracellular signaling pathways that regulate renin synthesis and secretion^46^.

In conclusion, our study provides a new understanding of the Ca^2+^ dynamics that underlie AngII effects on renin secretion in native JGCCs. Circulating renin levels result from a balance between the stimulation and inhibition of renin secretion, both essential to maintaining appropriate renin levels that sustain normal BP and fluid-electrolyte balance. Overall, this study highlights the often-overlooked inhibitory step whereby the end product of the renin-angiotensin cascade, AngII, exerts negative feedback on renin release through a complex regulation of [Ca^2+^]_int_ and, in doing so, maintains homeostasis. Understanding the nature of Ca^2+^ dynamics in JG cells could lead to the development of novel and safer therapeutic approaches for cardiovascular and renal diseases.

## Acknowledgments

We thank Thomas Wagamon for their expert animal care.

## Nonstandard Abbreviations and Acronyms

AngII: Angiotensin II
BP: blood pressure
cAMP: cyclic adenosine monophosphate
CPA: cyclopiazonic acid
Cx: Connexin
EGTA: ethyl-glycol tetraacetic acid
ER: endoplasmic reticulum
GJ: gap junction
[Ca^2+^]_int_: intracellular calcium
IP3: inositol 1,4,5-trisphosphate
IP3R: inositol 1,4,5-trisphosphate receptor
JG: juxtaglomerular
JGA: juxtaglomerular apparatus
JGCC: juxtaglomerular cell cluster
ROI: region of interest
SERCA: sarcoplasmic/endoplasmic reticulum calcium-ATPase
SOCC: store-operated calcium channels
SOCE: store-operated calcium entry
STIM: calcium-sensing stromal interaction molecule
TRP: transient receptor potential
VDCC: voltage-dependent calcium channels
VSMC: vascular smooth muscle cell

## Supplemental Methods

### Animals

*Ren1^c-Cre^* mice, where Cre expression is under the control of the *Ren1^c^* promoter^15^, were bred to B6J.Cg-*Gt(ROSA)26Sor^tm95.1(CAG-GCaMP6f)Hze^/Mwar^J16^* (Jackson Laboratory, Bar Harbor, ME, Strain #028865) generating mice with JG cell-specific GCaMP6 expression (*Ren1^c^-GCaMP6f* mice; *GCaMP6f^fl/+^; Ren1^c-Cre/+^*). *Ren1^c^-GCaMP6f* male and female mice between 30 and 100 days of age were used for each experiment except where specifically noted. C57BL/6J mice were obtained from the Jackson Laboratory (Strain #000664). The Major Resources Table details the number of animals used, their ages, sexes, and the groups compared in each experiment.

Genotyping of the mice was performed by TransnetYX (Cordova, TN). All animals utilized in this study were maintained on a C57BL/6J background. Mice were housed in groups in a room with controlled temperature and humidity, following a 12-hour light/dark cycle. All animals were handled in accordance with the National Institutes of Health guidelines for the care and use of experimental animals. The Institutional Animal Care and Use Committee at the University of Virginia approved the study.

### Kidney slice preparation

*Ren1^c^-GCaMP6f* mice were anesthetized with an intraperitoneal injection of tribromoethanol (300 mg/kg), and then the right kidney was dissected. Kidneys were sectioned (75 μm-thick) in ice-cold PIPES incubation buffer (20 mM PIPES, 119 mM NaCl, 4 mM KCl, 2 mM CaCl_2_, 1 mM MgCl_2_, 25 mM D-Glucose, 5 mM NaHCO_3_, adjusted pH 7.25–7.3 with 10 N NaOH to set Na 154 mM) using Leica VT1200 S fully automated vibrating-blade microtome (Leica, Wetzlar, Germany). Sections were kept at 37 °C for 20 min and then allowed to return to room temperature for the duration of the experiment. PIPES Imaging buffer (20 mM PIPES, 104 mM NaCl, 4 mM KCl, 2 mM CaCl_2_, 1 mM MgCl_2_, 25 mM D-Glucose, 0.1% bovine serum albumin, adjusted pH 7.25–7.3 with 10 N NaOH to set Na 134 mM and 280 mOsm) included with drugs were perfused at a rate of 2.5 mL/min and 37 °C throughout acquisition. The following drugs were used: Angiotensin II (50 pM–300 nM, Thermo Fisher Scientific, Waltham, MA), Nifedipine (10 μM, Sigma-Aldrich, St. Louis, MO), TTA-P2 (10 μM, Merck Millipore, Billerica, MA), SAR7334^26^ (10 μM, MedChemExpress, Monmouth Junction, NJ), Pico145^25^ (10 μM, MedChemExpress), Ruthenium Red (10 μM, Sigma-Aldrich), Cyclopiazonic acid (20 μM, Cayman Chemicals, Ann Arbor, MI), 2-Aminoethoxydiphenyl borate (100 μM, Thermo Fisher Scientific), Dantrolene (10 μM, MedChemExpress), Gadolinium (5–50 μM, MedChemExpress), GSK-7975A^32^ (10 μM, MedChemExpress), and Carbenoxolone (100 μM, Thermo Fisher Scientific).

### *Ex vivo* Ca^2+^ image acquisition

Kidney slices were imaged as previously described with minor modification^7^. Briefly, kidney slices (75 μm-thick) were imaged in a perfusion buffer on a widefield fluorescence microscope (Zeiss Axio-Examiner, Carl Zeiss Meditec, Dublin, CA) equipped with a 63x water-dipping objective lens, X-Cite XLED light source (Excelitas Technologies Corp., Waltham, MA), and a PM-7D magnetic stage with RC-27L large bath chamber (Warner Instrument Corp., Hamden, CT) configured for a perfusion system. Images were acquired at 10 Hz with an sCMOS camera (Hamamatsu Orca-Flash 4.0) using Slidebook 6 software (Intelligent Imaging Innovations, Denver, CO). Sections were pretreated for ∼20 min with Angiotensin II (50 pM–300 nM) before the start of each experiment. For most experiments, a baseline (AngII-pretreated) 10-minute acquisition period was followed by a 15-minute perfusion of hormone/drug. Slidebook files were converted to high-quality multipage TIFFs for analysis using ImageJ/Fiji.

### *In vivo* Ca^2+^ image acquisition

Under continuous anesthesia with a 2.0% isoflurane inhalant administered via nose-cone (VetEquip V-1 Table Top Anesthesia system, VetEquip Inc., Pleasanton, CA), animals were prepared by removing their hair and cleansing their skin with 70% ethanol. Saline-filled PE10 polyethylene catheters (Becton Dickinson; 0.28 mm internal diameter) were inserted into the abdominal cavity through an 18-gauge needle for AngII intraperitoneal injection. The prepared mouse was then positioned prone on a warming pad to maintain body temperature. Once positioned, a 1.0–1.5 cm incision on the left dorsal surface exposes the left kidney. A coverslip-bottom surgical window (130–170 μm-thick) with a 0.625-inch diameter round metal frame was attached to the kidney by gluing the rubber rings surrounding the kidney surface to the round frame plate and the skin with a portion of the kidney surface. Then, the surgical windows were immobilized using a custom-made vise by clamping the metal plate attached to the imaging window. Mice were placed on the stage for kidney multiphoton microscopy imaging, and deionized-distilled H_2_O was poured above the coverslip-bottom surgical window for objective immersion. Images were acquired using a Bruker Ultima 2Pplus multiphoton microscope (Bruker, Billerica, MA) with a Nikon 16× water immersion objective, numerical aperture = 0.8, (MRP07220cfx, Nikon Instruments Inc., Melville, NY), powered by an Axon 920 nm laser (2261788, Coherent Corp., Palo Alto, CA) and a GaAsP photocathode photosensor module with photomultiplier tube detectors (H16201P-40-S1, Hamamatsu), emitting at 300–740 nm with peak detection at 520 nm for GCaMP6f. The potential toxicity of laser excitation and fluorescence to the cells was minimized through low laser power and high scan speeds, ensuring that total laser exposure was kept to a minimum. Fluorescence images were collected in volume or time series (29.989 Hz) and converted to high-quality multipage TIFFs using PrairieView image acquisition software (Bruker, version 5.8.64.800). An electro-tunable lens (Optotune EL-10-30-TC) was used to capture high-speed volume z-series. The JG cluster exhibiting the highest activity within the visible range was selected for analysis. AngII (100 ng/g) suspended in 0.3 mL saline was injected within a minute through the polyethylene catheters while applying the laser and capturing the images. To minimize the effects of anesthesia, surgery, and laser-induced invasion on fluorescence and circulation, imaging was completed within 20 minutes from setting the imaging windows for each mouse.

### Signal extraction and analysis

Each multipage TIFF time series from *ex vivo* and *in vivo* Ca^2+^ image acquisition was cropped and analyzed using the Mesmerize platform^17^ and custom MATLAB software. Imaging data were motion-corrected using the CaImAn tool^18^, and then ROIs were detected with CNMF(E)^47^ under the Mesmerize analysis platform. Ca^2+^ transient spikes in each time series are identified using custom MATLAB software, which combines an automated spike detection algorithm with manual corrections for erroneous or missing spikes. Ca^2+^ activity is presented in raster plots and quantified to determine mean spikes/cell, mean frequency/cell, and bursting activity. To assess manipulation-dependent changes in cellular activity, Ca^2+^ spikes are organized in the 1-minute bin, and mean spikes/cells are averaged across ROIs within a kidney slice or native kidney. To quantify the changes in fluorescent intensity, a hand-drawn ROI was placed closely over the JGCC, and background fluorescence was subtracted from the fluorescent intensity in each serial image. The fluorescent intensities were normalized to baseline periods to measure the changes as GCaMP6f F/F_0_ values.

### Burst detection

Spikes were grouped into bursts, as we previously reported^7^. Briefly, all inter-spike period distribution was fitted with a mixture of Gaussian and exponential distributions. All of the inter-spike period values at the intersection of the Gaussian and exponential components of the distribution were taken as a threshold to determine the maximum inter-spike periods within a burst (see also Figure S3). Bursts were defined as three or more consecutive Ca^2+^ spikes within a calculated time threshold from the previous spike. Burst properties (e.g., the number of bursts, spike number, duration) were averaged across individual cells to derive a mean value per kidney.

### Plasma renin ELISA

For basal plasma renin measurement, blood was collected from animals anesthetized with an intraperitoneal injection of tribromoethanol (300 mg/kg). Blood samples were placed in tubes containing EDTA or heparinized plasma separator tubes (Becton Dickinson, Franklin Lakes, NJ). Plasma samples were collected from blood following centrifugation at 1,000 g for 20 minutes at 4 °C. Renin concentration was determined using ELISA (RayBiotech, Norcross, GA)^3,8^.

### Kidney slice renin secretion assay

Kidney sections from male C57BL/6J mice (60–89 days old) were cut into 75 μm-thick slices as described in the kidney slice preparation section. In a 24-well plate, each kidney section was placed in 1.0 mL of PIPES imaging buffer. After a 15–30 minute incubation (Baseline, same incubation time per assay), a 1 mL sample of the solution was collected, and the sections were moved to a new 24-well plate containing fresh media with various concentrations of AngII or drugs. After an additional 30 minutes, a final sample (Treatment) was obtained. The renin concentration was determined using ELISA following the manufacturer’s instructions (RayBiotech). To account for variations in the number of JG cells in each kidney slice, the renin concentration of Treatment samples was divided by that of each Baseline sample, which was calculated as Fold renin secretion.

### Carotid arterial pressure measurement and acute *in vivo* renin secretion assay

*Ren1^c^-GCaMP6f* mice were anesthetized with 2.0% isoflurane inhalation and maintained at 37.5°C on a heating pad. Heparinized saline-filled polyethylene catheters (PE10, Becton Dickinson; 0.28 mm internal diameter) were inserted into the right carotid artery using an RX104A transducer connected to a data acquisition system and AcqKnowledge software (Biopac Systems, Inc., Goleta, CA) to record the carotid arterial pressure continuously^8^. After recording pressure for 10 minutes as baseline blood pressure, 50 μL of blood was collected from the right jugular vein to measure the plasma renin level before the treatment. 10 minutes later, continuous intraperitoneal administration of AngII or an equivalent volume of saline was initiated.

Subsequently, 50 μL of blood was collected from the right jugular vein 30 minutes later to measure the plasma renin level after the treatment. The following methods for plasma renin ELISA were the same as those in basal plasma renin measurement.

### Immunohistochemistry**/**Immunofluorescence staining and histological analysis

Mice were anesthetized with intraperitoneal injection of tribromoethanol (300 mg/kg). Both side kidneys were overnight fixed in 4% paraformaldehyde (PFA), 10% formalin, or Bouin’s solution, and embedded in paraffin. For frozen block making, tissues were fixed in 4% PFA under vacuum for 2 hours at 4°C using a vacuum desiccator. After washing with phosphate-buffered saline (PBS) 2 times, the samples were immersed in 30% D-sucrose/PBS overnight at 4°C and frozen in O.C.T. (Thermo Fisher Scientific). For Periodic acid–Schiff (PAS) staining, Bouin’s fixed paraffin sections were stained with PAS reagent.

For immunofluorescence staining of renin, the frozen blocks were sectioned at 8 μm-thickness. After washing with PBS, the sections were post-fixed with 4% PFA for 30 minutes. After the blocking with 3% BSA, 5% donkey serum, and 0.04 % cold fish skin gelatin, sections were incubated with the rabbit monoclonal anti-renin antibody (1:5,000, Abcam, Cambridge, MA, ab212197) at 4 °C overnight. After the washing, sections were incubated with Alexa Flour 647 donkey anti-rabbit IgG (1:400, Vector Laboratories, Burlingame, CA, BA-1000) at room temperature for 2 hours. After the washing, sections were incubated with a quencher (Vector TrueView Autofluorescence Quenching Kit #SP-8400, Vector Labs) for 5 min to decrease autofluorescence. Nuclei were stained with Hoechst 33342 (1:5,000, Thermo Fisher Scientific).

For Immunohistochemistry staining for renin and α-smooth muscle actin (α-SMA), tissue sections from paraffin blocks fixed with Bouin’s solution were deparaffinized, rehydrated, and treated with 0.3% hydrogen peroxide in methanol. After the blocking with 3% BSA and 2% goat serum or horse serum, sections were incubated with the rabbit polyclonal anti-renin antibody generated in our laboratory (1:500) or the anti-α-SMA antibody (1:10,000, Millipore-Sigma, Burlington, MA, A2547) at 4 °C overnight^3,8^. After the washing, sections were incubated with biotinylated secondary antibody, goat anti-rabbit IgG (1:200, Vector Laboratories, BA-1000) or horse anti-mouse IgG (1:200, Vector Laboratories, BA-2000) for renin or α-SMA, respectively, at room temperature for 30 minutes. Staining was amplified using the Vectastain ABC kit (Vector Laboratories) and developed with 3,3’-diaminobenzidine Substate (Vector Laboratories). The sections were counterstained with hematoxylin (Millipore-Sigma), dehydrated, and mounted with Cytoseal XYL (Thermo Fisher Scientific). Images were captured using a Zeiss Imager M2 microscope equipped with an Apotome2 with an AxioCam 506 mono and an AxioCam 305 color camera (Carl Zeiss, Oberkochen, Germany).

The JG index was determined as the percentage of renin-positive JG cells per total glomeruli^8^. We counted the number of renin-expressing cells in each JG cell and along the arterioles with visible glomeruli attached to determine the number of renin-expressing cells per section.

### RNA extraction and quantitative RT-PCR

Renal cortices were removed and placed in RNAlater Stabilization Solution (Thermo Fisher Scientific) overnight at 4°C, then stored at −20°C. RNA was extracted from the renal cortices using TRIzol reagent (Thermo Fisher Scientific) and purified using an RNeasy Mini Kit (Qiagen, Germantown, MD), following the manufacturer’s protocol. Reverse transcription (RT) was conducted with oligo(dT) primer (Promega, Madison, WI) and M-MLV Reverse Transcriptase (Promega), following the manufacturer’s instructions. Quantitative PCR was executed with SYBR Green I (Thermo Fisher Scientific) in a CFX Connect Real-Time PCR Detection System (Bio-Rad, Hercules, CA). PCR was performed using the following primers: *Ren1*, forward: 5’-ACAGTATCCCAACAGGAGAGACAAG-3’, reverse: 5’-GCACCCAGGACCCAGACA-3’; *Rps14*, forward: 5’-CAGGACCAAGACCCCTGGA-3’, reverse: 5’-ATCTTCATCCCAGAGCGAGC-3’. The mRNA expression was normalized to *Rps14* expression, and the changes in expression were determined by the ΔΔCt method and were reported as relative expression compared to control mice^3^.

### Bubble plot for single-cell RNA sequence

Our previous single-cell RNA sequencing data from embryonic and adult *FoxD1^cre/+^; R26R^mTmG^* mice kidneys (GSE218570^9^) was used for dot plot visualization of L-type and T-type VDCC α_1_-subunits, canonical TRP (TRPC) and vanilloid TRP (TRPV) subfamilies, ORAI/STIM complex, endoplasmic reticulum channels, and GJ protein genes set in renin-lineage cells. The dot plot was produced using custom versions of the functions from https://renin-analysis.readthedocs.io/en/latest/18.R_generate_bubble_plots/. The input to the calcBubble() function was the raw gene expression matrix. The input matrix was then log-normalized, and the z-score was transformed with the percentage of cells of each population expressing each gene calculated along with the z-scores for each gene across all cell populations in the gene expression matrix. The output from this function was then plotted using ggplot2.

### Statistical Analysis

Statistical analyses were performed using GraphPad Prism software (ver10.2.3) and R (ver4.4.1). The normal distribution of the multi-comparison and paired-comparison data was evaluated with the Shapiro–Wilk test, followed by a paired t-test, one-way ANOVA, or two-way ANOVA test for normally distributed data. Based on the test requirements, Mann-Whitney U test, paired t-test, one-way ANOVA with Tukey’s multi-comparison test, two-way ANOVA with Šídák’s multi-comparison test, and linear mixed model. Statistical tests and results are shown in Figure legends. P<0.05 was considered significant. Data are shown as mean±SEM. Schematics created with BioRender.com. The ridgeline plots were generated using ggplot2 libraries of R.

The hypotheses were tested using a linear mixed model that accounts for the within-subject correlation of pseudoreplicates. The multi-level analysis’s number of pseudoreplicates (ROIs/slice) and genuine replicates (mice) is reported in each figure legend where applicable. For Ca^2+^ imaging studies in which each mouse contributed exactly one slice per experimental condition, the mouse is considered a genuine replicate, while the slice and every ROI within the slice are considered pseudoreplicates. Therefore, the correlated error was modeled using a hierarchical structure, with mouse > kidney slice or native kidney > ROI.

---

**Figure S1.**
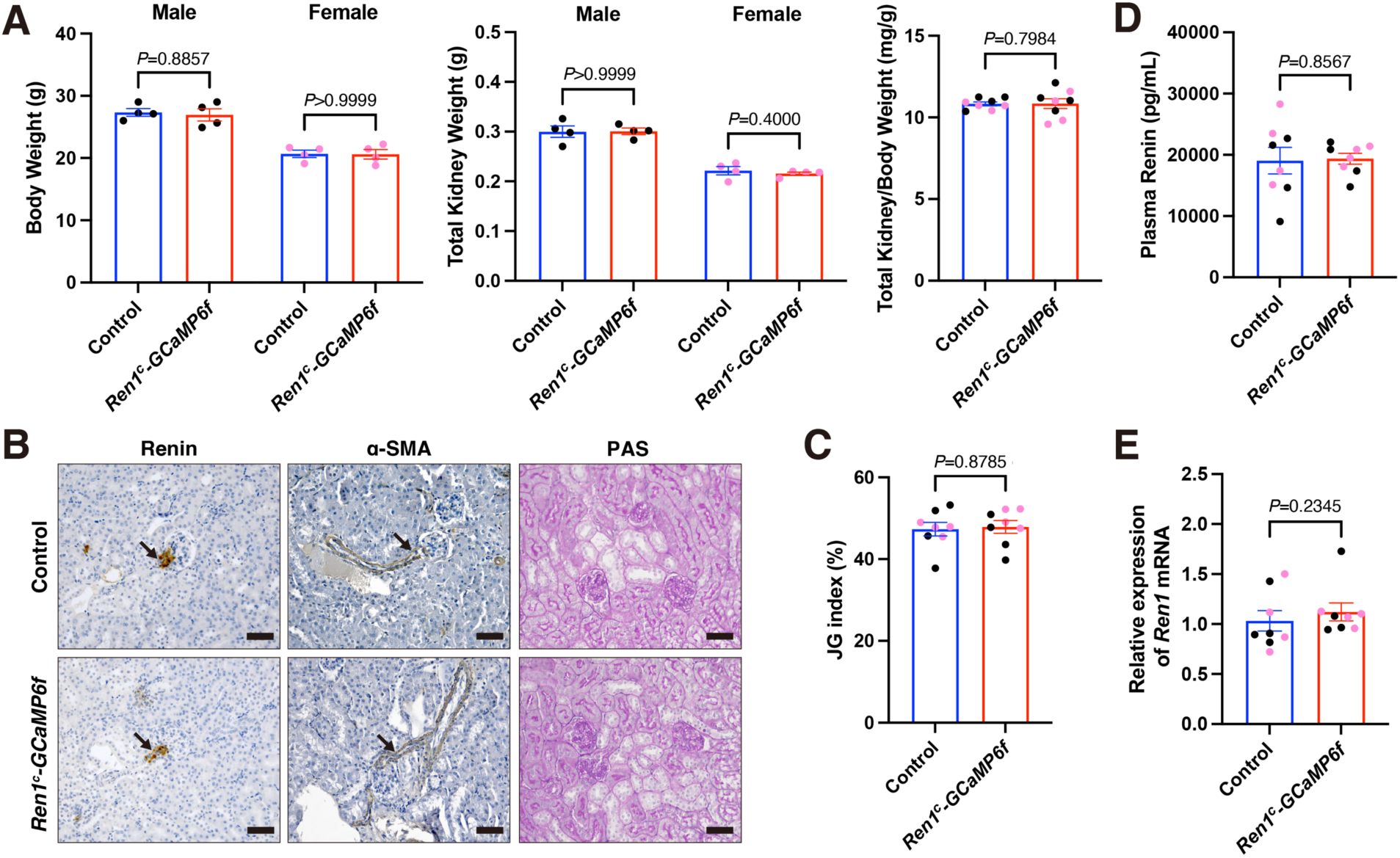
*Ren1^c^-GCaMP6f* mice show normal kidney development and renin expression in juxtaglomerular (JG) cells. **A,** Body weight and total kidney weight showed no significant difference between *Ren1^c^-GCaMP6f* mice (79–81 days old, n=4 for males and n=4 for females) and Control (81 days old of C57BL/6J mice, n=4 for males and n=4 for females) in both males and females (Mann-Whitney U test). The Total kidney/body weight ratio also showed no significant difference between *Ren1^c^-GCaMP6f* mice and Control mice (Mann-Whitney U test). **B,** Immunohistochemistry for renin and α-smooth muscle actin (α-SMA) and Periodic acid–Schiff (PAS) staining showed no abnormality of the kidneys from *Ren1^c^-GCaMP6f* mice. Brack arrows in the immunohistochemistry staining indicate the tips of afferent arterioles. Scale bar, 50 μm **C,** The quantification of renin staining JG area/glomerulus (JG index) showed no significant difference between *Ren1^c^-GCaMP6f* mice (Mann-Whitney U test). **D–E,** Plasma renin concentration (**D**) and kidney cortices *Ren1* mRNA (**E**) of *Ren1^c^-GCaMP6f* mice at normal states did not show significant differences compared to the Controls (Mann-Whitney U test). Black dots show male samples and purple dots show female samples. Data presented as mean±SEM.

**Figure S2.**
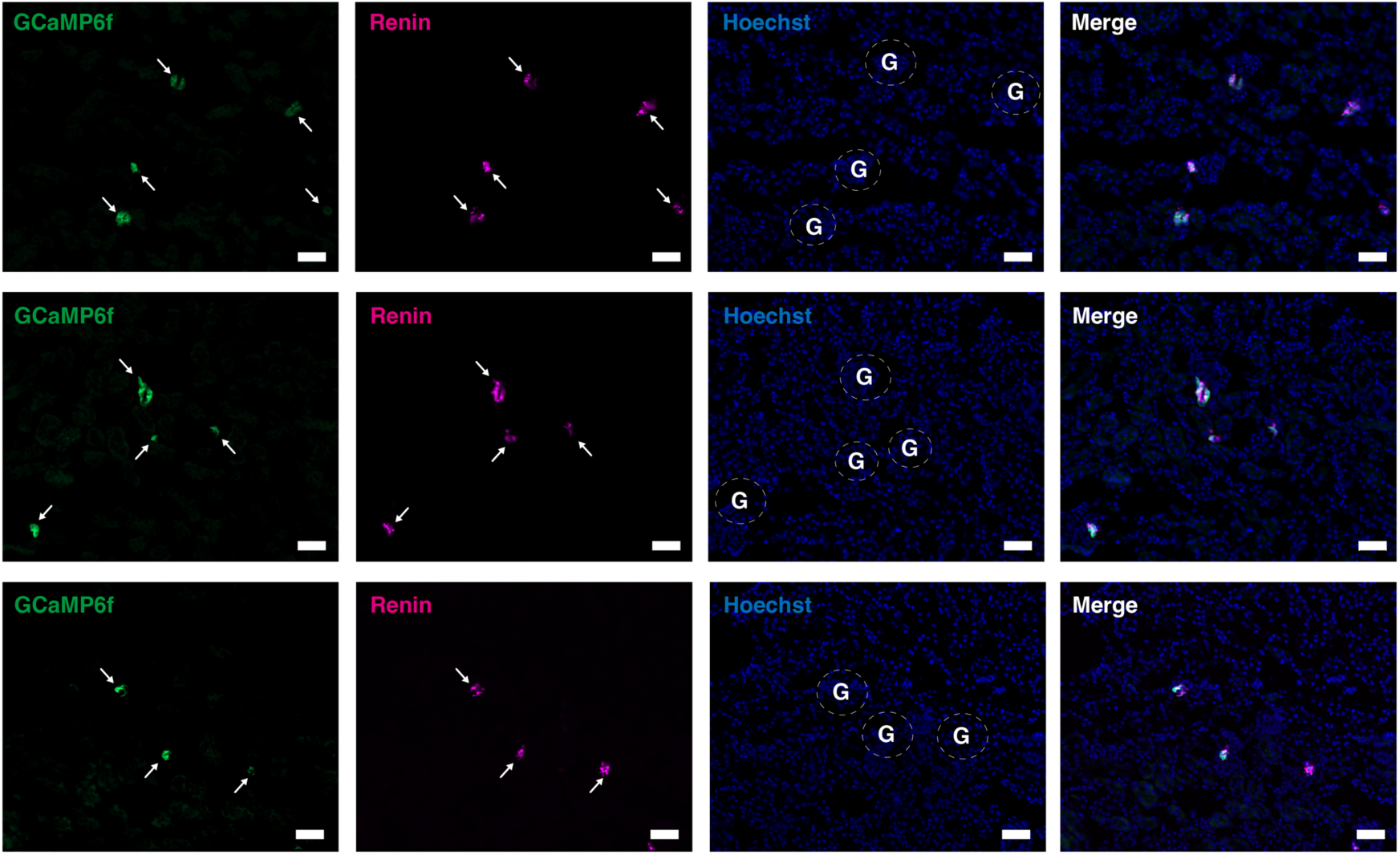
Colocalization of GCaMP6f and renin around the juxtaglomerular (JG) area of *Ren1^c^-GCaMP6f* mouse kidney. Immunofluorescence staining for renin (magenta) in the kidney cortex of the *Ren1^c^-GCaMP6f* mice. Nuclei are stained with Hoechst (blue). The merged image shows GCaMP6f (green) expressed in the JG area around glomeruli stained with an anti-renin antibody. White arrows point to the location for renin and GCaMP6f expressions. Dashed white circles indicate glomeruli. G; Glomerulus. Scale bar, 50 μm.

**Figure S3.**
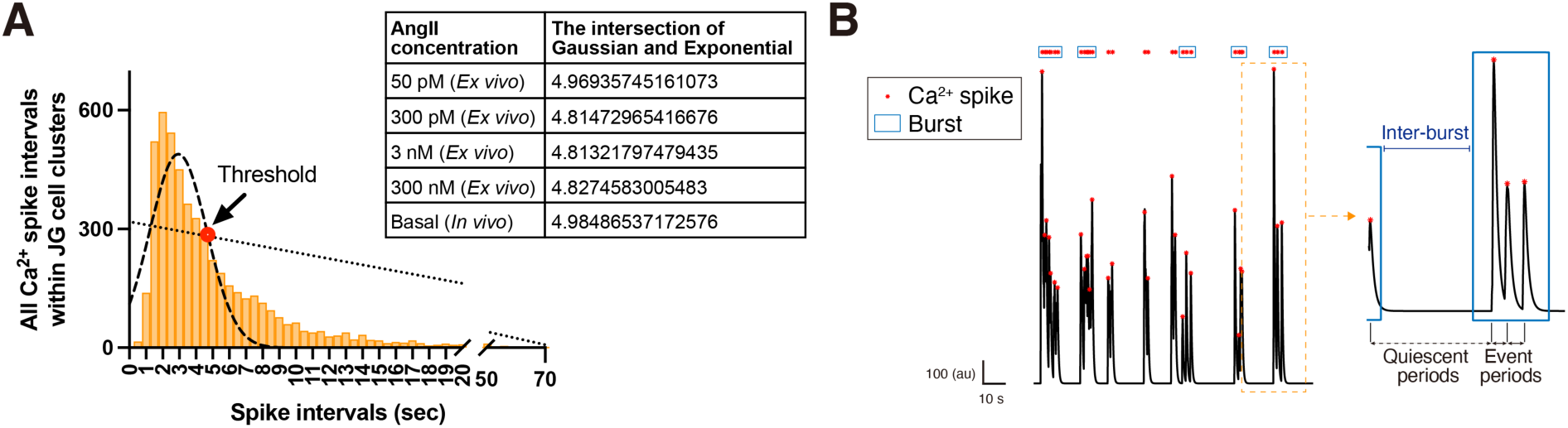
Calculations for determining threshold values for defining event periods. **A,** The histogram of all inter-spike periods across all regions of interest (ROIs) in five juxtaglomerular cell clusters (JGCCs) treated with 3 nM angiotensin II (AngII) is shown. The solid line indicates the Gaussian curve, and the dotted line indicates the exponential components. The intersection of the Gaussian and exponential components (red dot) is used for the Gaussian-exponential model (thresholds). The table shows thresholds for the maximum intra-burst interval as calculated from the Gaussian-exponential model for each AngII dose (50 pM, 300 pM, 3 nM, and 300 nM) in *ex vivo* Ca^2+^ imaging and basal state in *in vivo* Ca^2+^ imaging. **B,** Example of trace data and the successive Ca^2+^ spike (red dots) with burst assignment (blue squares). Burst requires three consecutive spikes with short intervals below the threshold (event periods), while spike intervals exceeding the threshold (quiescent period) indicate inactive inter-burst periods.

**Figure S4.**
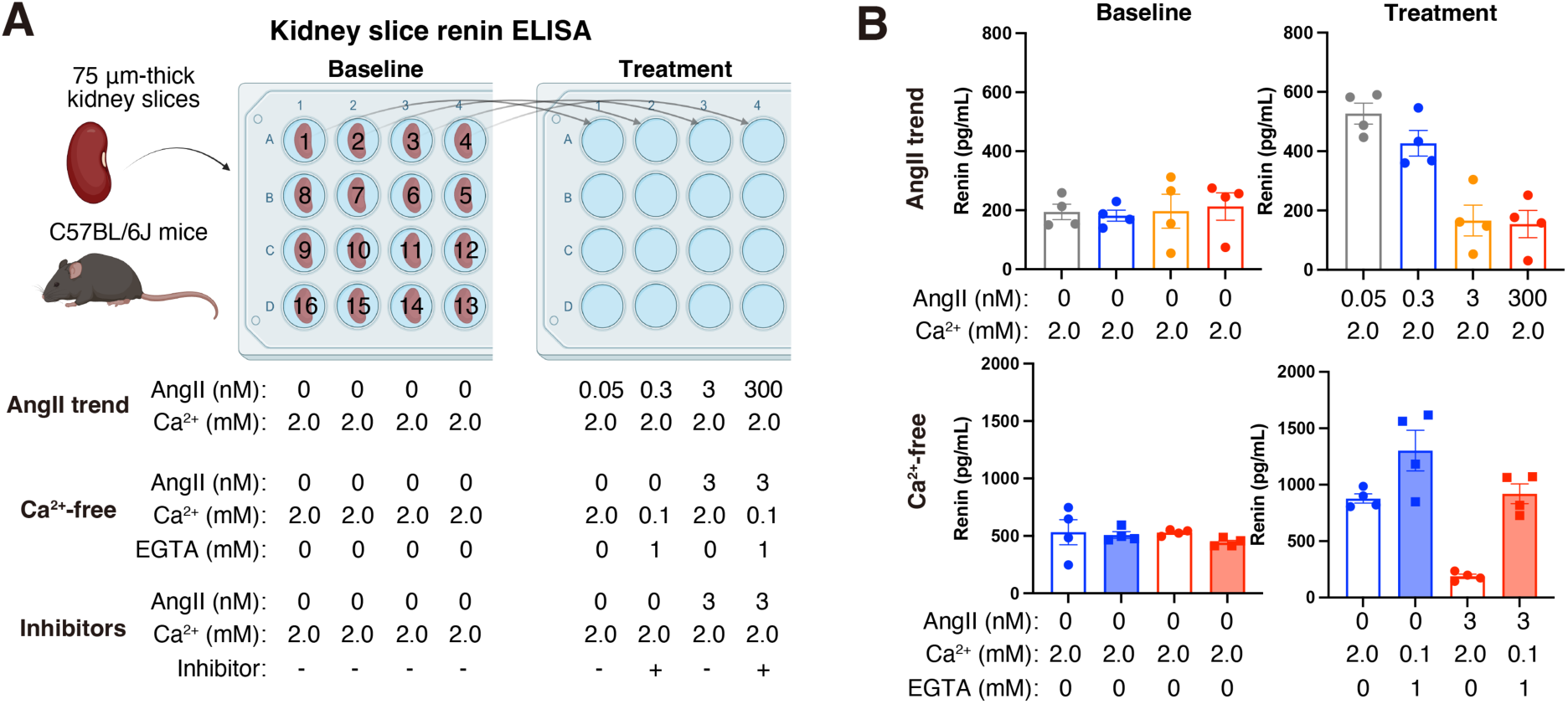
Overview of kidney slice renin ELISA. **A,** Schematic of kidney slice renin ELISA. A kidney from male C57BL/6J mice (60–89 days old) lying on its side was cut into 75 μm-thick slices from the topside of the kidney. Each kidney section was placed in 1.0 mL of PIPES imaging buffer with 2 mM Ca^2+^. To minimize the variation in the size of kidney sections across the groups, the sections were arranged according to the number illustrated on the 24-well plate. Following a 15–30 minute incubation (Baseline, with consistent incubation time for each assay), a 1 mL sample of the solution was taken. Then, the sections were transferred to a new well plate filled with 1 mL fresh media containing various angiotensin II (AngII) concentrations (AngII trend), ethyl-glycol tetraacetic acid (EGTA) with Ca^2+^ (Ca^2+^-free), and inhibitors shown below the scheme. After an additional 30 minutes, a final sample (Treatment) was obtained. ELISA determined the renin concentration in the samples. **B,** Renin concentration in samples of Baseline and Treatment with AngII trend and Ca^2+^-free. To address differences in the number of juxtaglomerular (JG) cells per kidney slice, the renin concentration of Treatment samples was calculated as a ratio to each Baseline sample, which was referred to as fold renin secretion (AngII trend; Figure 2F, Ca^2+^-free; Figure 3B). The same strategy was employed in treatment with Gd^3+^ and GSK-7975A (Figure 3F) and carbenoxolone (Figure 5D). Data presented as mean±SEM.

**Figure S5.**
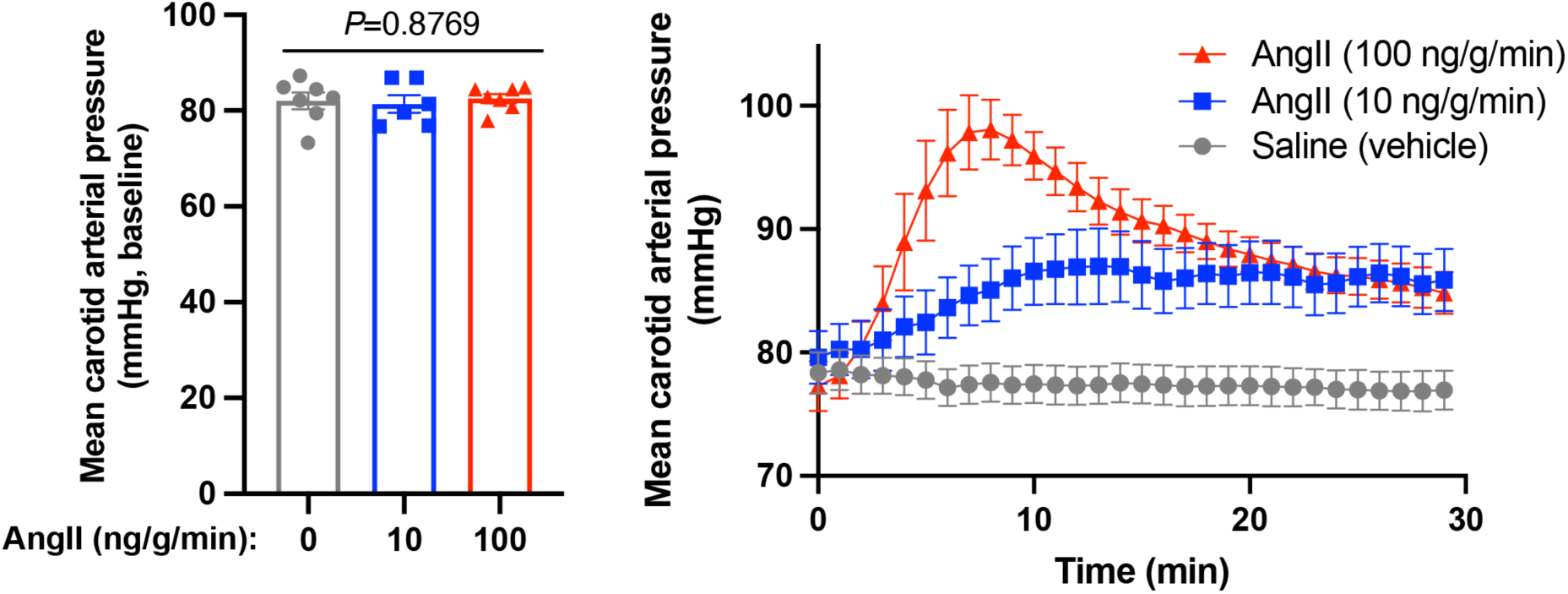
Carotid arterial pressure data in acute *in vivo* renin secretion assay. Mean carotid arterial pressure at baseline for 10 minutes (left, One-way ANOVA) and time-course of mean carotid arterial pressure over 30 minutes under intraperitoneal administration with saline (vehicle), 10 ng/g/min, and 100 ng/g/min angiotensin II (AngII, n=7, 6, and 7 *Ren1^c^-GCaMP6f* mice, respectively). Initially, there was no difference in carotid arterial pressure among the group. Following the start of treatment, AngII significantly increased the carotid arterial pressure in a dose-dependent manner, both earlier and to a greater extent, while the vehicle had minimal impact. Data presented as mean±SEM.

**Video S1. Angiotensin II (AngII) elicits robust oscillatory Ca^2+^ signals coordinated within the juxtaglomerular cell cluster (JGCC).**

Time series of captured JG cell images at the JG area in a kidney slice from *Ren1^c^-GCaMP6f* mice under 3 nM AngII containing 2 mM Ca^2+^ buffer perfusion. Yellow arrows point to the JGCC during intense GCaMP6f expression. Scale bar, 20 μm.

**Video S2. A pan gap junction (GJ) inhibitor suppresses angiotensin II (AngII)-elicited Ca^2+^ oscillations within the juxtaglomerular cell cluster (JGCC).**

Time series of captured JG cell images at the JG area in a kidney slice from *Ren1^c^-GCaMP6f* mice under 3 nM AngII with (left) and without (right) 100 μM carbenoxolone (CBX) treatment. Yellow arrows point to the JGCC during intense GCaMP6f expression Scale bar, 20 μm.

**Video S3. Intravital imaging of *Ren1^c^-GCaMP6f* mouse kidney reveals robust oscillatory Ca^2+^ signal coordinated within the juxtaglomerular cell cluster (JGCC) under physiological conditions.**

Captured fluorescence volume series of GCaMP6f at the JGCC (29.989 Hz, 400 multi-page TIFF). Yellow arrows point to the JGCC during intense GCaMP6f expression. Scale bar, 20 μm.

**Video S4. Continuous intravital imaging of juxtaglomerular cell cluster (JGCC) in *Ren1^c^-GCaMP6f* mouse before and after angiotensin II (AngII) administration.**

Time series of GCaMP6f signals from the JGCCs in *Ren1^c^-GCaMP6f* mice before and after AngII intraperitoneal administration (left) and the corresponding GCaMP6f F/F_0_ (right). Yellow arrows point to the JGCC during intense GCaMP6f expression. Scale bar, 20 μm.

Data are provided separately.

